# Evolution of Insulin, Insulin-like Growth Factor, and Their Cognate Receptors in Vertebrates, Invertebrates, and Viruses

**DOI:** 10.1101/2025.07.03.661983

**Authors:** Martina Chrudinová, Jeffrey M DaCosta, Dogus Dogru, Ruixu Huang, Robert Reiners, Pierre De Meyts, Emrah Altindis

## Abstract

The insulin and insulin-like growth factor (IGF) system regulates essential biological functions such as growth, metabolism, and development. While its physiological roles are well characterized, the evolutionary origins and molecular diversification of its ligands and receptors remain incompletely defined. Here, we present the most comprehensive phylogenetic and sequence conservation analysis of this system to date, using over 1,000 sequences from vertebrates, invertebrates, and viruses. Our analyses reveal that insulin, IGF-1, and IGF-2 form distinct monophyletic clades that diverged after the emergence of vertebrates, with IGF-1 being the most conserved ligand. We show that IGF1R-binding residues, especially in the A- and B-domains of IGF-1, are highly conserved across vertebrates, while insulin’s Site 2 residues, which overlap with its dimerization and hexamerization surface, are more variable—correlating with the loss of hexamer formation in hystricomorphs, reptiles, and jawless fish. Unexpectedly, we identify a 12–amino acid insert in the insulin receptor (IR) of turtles and tortoises, previously thought to be unique to mammalian IR-B isoform, suggesting an earlier evolutionary origin of isoform diversity. We also show that marsupials and monotremes retain ancestral receptor domain features shared with reptiles and birds, and that avian insulins, particularly A-chain residues, are unusually conserved. Viral insulin/IGF-like peptides (VILPs) fall into two distinct clades that resemble either IGFs or insulin. Together, these findings illuminate the evolutionary architecture of the insulin/IGF system, highlight unexpected lineage-specific adaptations, and provide a framework for understanding hormone-receptor function across biology and therapeutic design.

## Introduction

The insulin/insulin-like growth factor (IGF) system is a complex regulatory network of hormones, receptors, and binding proteins that collectively regulate a broad spectrum of biological processes, including metabolism, growth, and development. In vertebrates, this system comprises three peptide hormones: insulin and two insulin-like growth factors (IGF-1 and IGF-2) [1,2]. Insulin primarily serves as a central regulator of glucose homeostasis [3], while IGFs primarily function as growth factors. IGF-1 promotes cellular growth, proliferation, and differentiation, whereas IGF-2 plays a critical role in embryonic development and has additionally been implicated in neuroprotection, and memory formation/consolidation [1,4].

These ligands exert their effects through binding to and activating their cognate receptors, including the insulin receptor (IR) and IGF-1 receptor (IGF1R), both of which are members of the receptor tyrosine kinase family [1,5,6].

Affirming the evolutionary conservation of the insulin/IGF system across species, insulin-like peptides (ILPs) have also been identified in invertebrates. In contrast to vertebrate organisms, which typically possess a single copy of insulin, IGF-1, and IGF-2, research on invertebrate model organisms has shown that they frequently express a multitude of ILPs. For instance, *Drosophila melanogaster* harbors eight ILPs, known as *Drosophila* insulin-like peptides (DILPs) [7], and *Caenorhabditis elegans* encodes 40 ILPs [8]. ILPs have been shown to be involved in numerous functions, including stress response, lifespan, longevity, and sex-related functions [9–11]. Interestingly, despite their diversity, most invertebrate species mediate the effects of multiple ILPs through a receptor encoded by a single gene [12–14], even though receptor duplication has been observed in some species [15,16].

Despite their evolutionary distance, vertebrate insulin/IGFs and invertebrate ILPs share highly conserved primary and tertiary structures. Vertebrate insulin is a double-chain peptide composed of an A-chain and a B-chain. In proinsulin, which represents the immature form of insulin, a connecting peptide (C-peptide) links the two chains, but it is removed during proinsulin processing. In mature insulin, the separate A- and B-chains are held together by two disulfide bonds, with one extra disulphide bond present within the A-chain [17–19]. In contrast, vertebrate IGFs are single-chain peptides comprising A-, B-, C- and a short D-domain. The C-domain remains present in mature IGFs and connects the B- and A-domains. An analogous disulphide bonding pattern to those in insulin is also present in IGFs, and therefore serves as a key motif characterizing the insulin/IGF family members in vertebrates [19–21]. The A- and B-chains of insulin and the A- and B-domains of IGFs exhibit high sequence conservation within species. For instance, in humans, there is approximately 50% amino acid sequence conservation between insulin and IGFs, and about 70% between IGF-1 and IGF-2 (Fig 1A). Most of invertebrate ILPs have a double-chain structure similar to vertebrate insulin, although some peptides that retain an IGF-like single-chain structure have been identified [14,19,22–27]. The cysteine residues critical for disulphide bond formation are highly conserved in ILPs, and most invertebrate peptides exhibit a disulphide bonding pattern similar to that of vertebrate peptides, although some variants with slightly modified patterns have also been reported [14,19,24,28–30].

**Fig 1.**
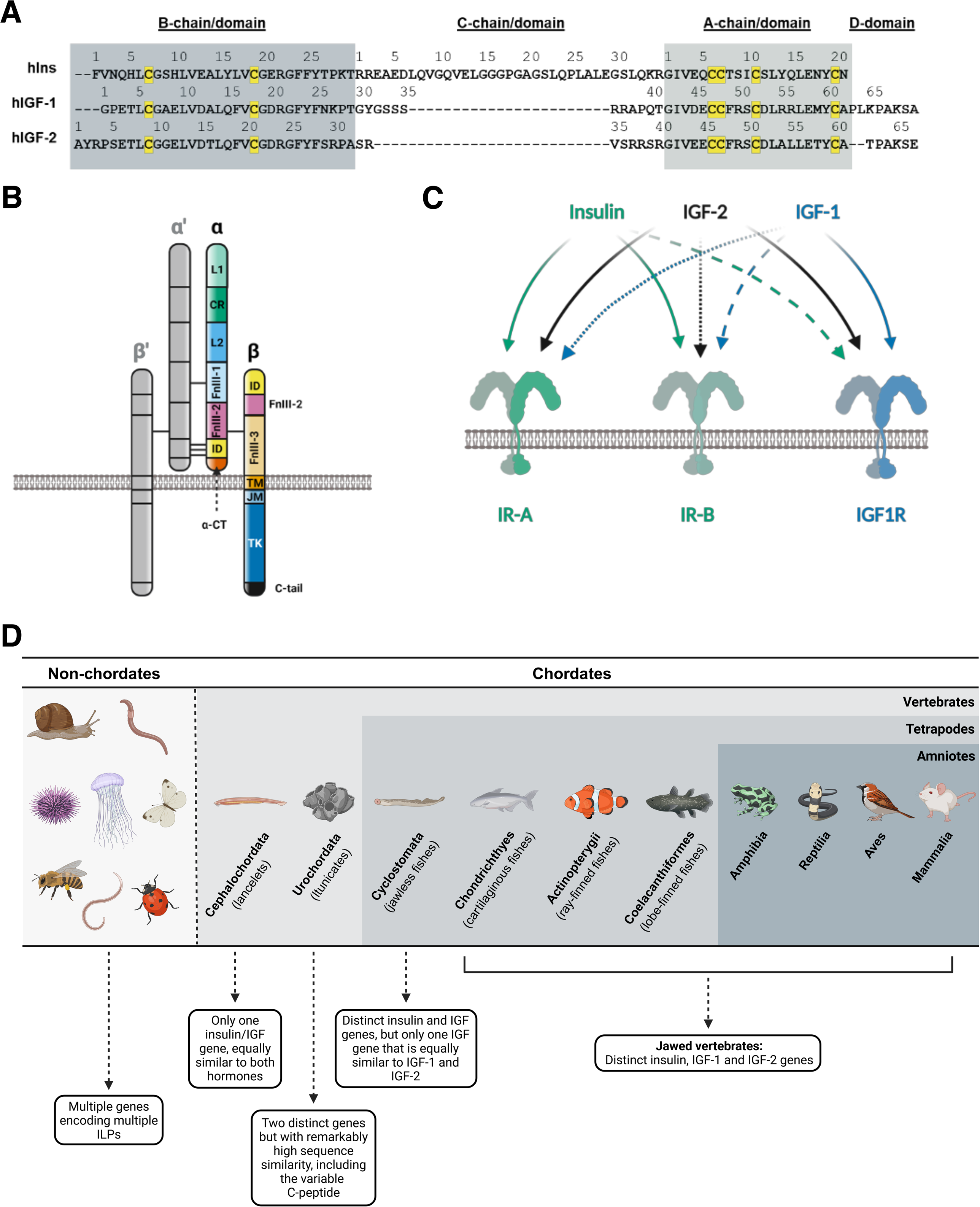
Overview of insulin, insulin growth factor (IGF), and insulin-like ligands and their cognate receptors. (A) Alignment of human insulin (hIns), IGF-1 (hIGF-1), and IGF-2 (hIGF-2) sequences. (B) Subunit and domain composition of insulin receptor (IR) and IGF-1 receptor (IGF1R). L1 and L2 - leucine rich domain 1 and 2; CR - cysteine rich region; FnIII-1, FnIII-2 and FnIII-3 - fibronectin type III domain 1, 2 and 3; ID - insert domain, TM - transmembrane domain; JM - juxtamembrane domain, TK - tyrosine kinase domain; α-CT - C-terminal region of the α-subunit; C-tail - C-terminal region of the β-subunit. (C) Schematic representation of ligand- receptor binding in the insulin/IGF system. (D) Evolution of insulin, IGF, and insulin-like receptors in animals.

Vertebrate IR and IGF1R are disulfide-linked heterotetrametric transmembrane receptors that share the same domain composition, each with two α (α and α’) and two β (β and β’) subunits. The α-subunits are extracellular and contain the ligand binding sites, whereas the β- subunits span the membrane and contain the catalytically active tyrosine kinase domain intracellularly (Fig 1B). In mammals, IR can be expressed as two isoforms (IR-A and IR-B), with IR-A lacking 12 amino acids in the C-terminal region of the α-subunit (α-CT) due to alternative splicing of the IR gene exon 11 [5]. IR-B is primarily expressed in insulin-sensitive tissues (e.g., liver, adipose tissue, skeletal muscle) in mammals, whereas IR-A is found in additional tissues like brain, heart, pancreas, spleen, and fetal tissue [31–33]. Although each vertebrate ligand can bind to its own receptor with high affinity, significant cross-talk occurs due to structural similarities between ligands and receptors (Fig 1C). For instance, human insulin binds primarily to human IR (hIR) but can also bind to human IGF1R (hIGF1R), albeit with lower affinity. Human IGF-1 exhibits the opposite pattern. Human IGF-2, which lacks its own tyrosine kinase receptor, binds with a relatively high affinity to both hIR-A and hIGF1R [34,35]. Until recently, the consensus based on mutagenesis and structural studies was that insulin and IGF-1 bind to IR and IGF1R by crosslinking two binding sites on the receptors: “Site 1” (high affinity binding site) and “Site 2” (low affinity binding site) [36–41]. Recent cryoEM studies showed that up to four insulin molecules can bind to IR, unveiling a new binding sub-site [42,43]. This led to a revised nomenclature, with previously identified binding sites now labeled as sub-sites “Site 1a” and “Site 1b,” and the newly discovered site named “Site 2.” These sites are located within FnIII-1, α-CT, and L1 domains on IR, respectively. This manuscript adopts this nomenclature when discussing conservation in sites involved in IR binding. On the other hand, IGF1R was shown to only bind up to two IGF-1 molecules via two binding sites analogic to IR site 1a and 1b, but no binding site 2 has been identified. However, the CR domain was also shown to be involved in binding of IGF-1 to IGF1R in addition to FnIII-1, α-CT, and L1 domains [44].

The recent discovery of viral insulin/IGF-1 like peptides (VILPs) in six Iridoviridae viruses further expanded the insulin/IGF/ILP superfamily [45]. VILPs share 30-50% amino acid sequence homology with human insulin/IGFs, including the conservation of all six cysteines that are essential for insulin/IGF tertiary structure. Additionally, both in vivo and in vitro studies confirmed that the VILPs are active ligands of the insulin/IGF system [35,45–48]. This discovery prompted us to investigate the phylogenetic relationships among all insulin-like molecules, including VILPs, and their cognate receptors in order to update our current understanding [1–5] of the evolution of this superfamily across species (Fig 1D).

In this study, we tapped the expanding genomic and proteomic databases to broadly sample the available sequences of vertebrate insulins/IGFs, invertebrate ILPs, VILPs, and their receptors, and analyzed their evolution using phylogenetic and motif analyses.

## Results

### Phylogenetic analyses reveal key findings in relationships among ligand sequences

The phylogenetic analysis of the full ligand data set was completed using an alignment of 628 insulin, IGF-1, IGF-2, ILP, and VILP sequences. The maximum likelihood topology of all ligands (Fig 2) was rooted with the two cnidarian sequences (fresh-water polyp *Hydra vulgaris* and coral *Stylophora pistillata*) since Cnidaria is the oldest and most distantly related extant lineage of animals in our data set. Early-branching clades in this phylogeny contain ILP sequences from invertebrates, although the long branches and low transfer bootstrap expectation (TBE) nodal support values in this part of the tree illustrate considerable divergence among sequences. Also, the ILP sequences did not form a monophyletic group; two ILP sequences from the lancelet (*Branchiostoma belcheri*), along with sequences from a thrip (*Frankliniella occidentalis*), termite (*Zootermopsis nevadensis*), and crustacean (*Daphnia pulex*) were more closely related to insulins/IGFs than to other ILPs (clade marked as Insecta+Chordata+Crustacea in Fig 2). Insulin, IGF-1, and IGF-2 sequences each form a monophyletic group with the TBE value for the basal node of each group exceeding 90%, indicating robust support for these relationships. IGF-1 and IGF-2 groups are sister to each other, suggesting that each evolved from duplication of an IGF-like protein in an ancient common ancestor [29,49,50]. However, there are a few non-vertebrate sequences that are positioned between the insulin and IGF-1/2 groups. The insulin sequences are most closely related to Lymphocystis disease virus LCDV-Sa and LCDV-4 VILPs, although there is considerable divergence between these insulins and VILP sequences.

**Fig 2.**
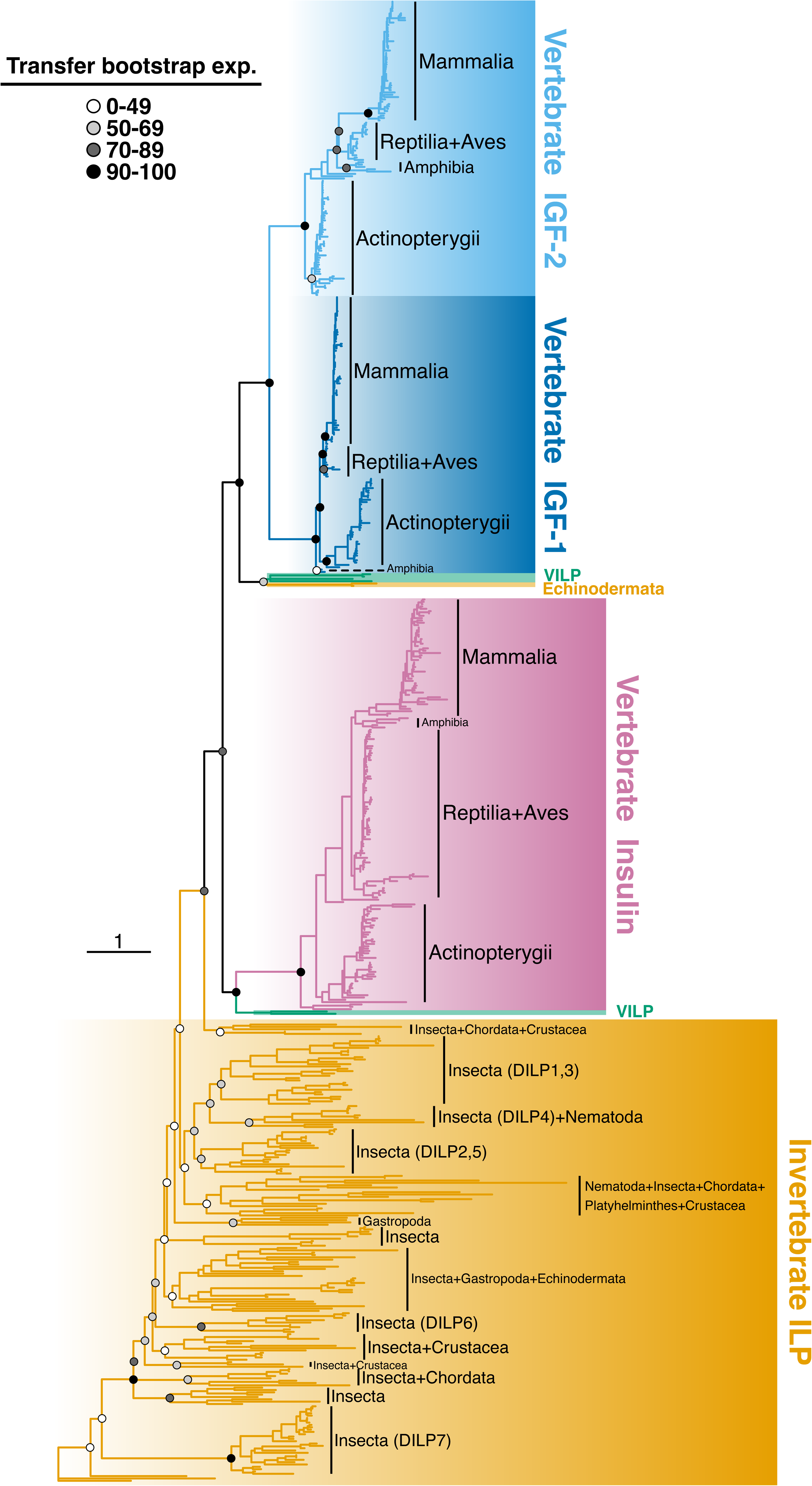
Phylogenetic relationships among animal insulin, insulin growth factor (IGF), and insulin- like ligands. The tree was generated using maximum likelihood based on an alignment of 628 sequences. Branches and clades are color-coded for different categories of ligands, and circles at key nodes illustrate clade support in the form of transfer bootstrap expectation (TBE) values. With all sequences included, this tree gives an overview of the relationships among sequences of insulin, IGF-1, IGF-2, invertebrate insulin-like proteins (ILPs), and viral insulin/IGF-like peptides (VILPs).

The IGF1/2 group is most closely related to two predicted insulin-like sequences from draft genomes of sea stars (Echinodermata; *Acanthaster planci* and *Patiria miniata*) and the remaining four VILP sequences (Mandarin fish Ranavirus, Grouper iridovirus, Singapore Grouper Iridovirus, and LCDV-1). Notably, there are long branches between these invertebrate/VILP sequences and the IGF/insulin groups, indicating significant evolutionary divergence between them. The branch lengths among VILP sequences are also long, indicating that they have a relatively ancient origin and also diverged from each other in the distant past. The placement of the two sea star sequences is curious as it is far from a third sequence from this taxonomic group (Echinodermata; sea cucumber *Stichopus japonicus*). These sea star sequences are predicted from draft genomes, and were only added to the dataset based on blastP searches for sequences similar to VILPs.

We performed a separate phylogenetic analysis of only ILP and VILP sequences (*N* = 205), which decreased overall divergence among sequences and generated a phylogeny with more robust nodal support values (Fig 3). The ILP+VILP data set was dominated by 70 *Drosophila* sequences. The VILP sequences cluster together (along with uncharacterized sea star proteins) with reduced sampling, but again long branches between sequences, particularly between LCDV-Sa and LCDV-4 from other VILPs, demonstrate that they are highly divergent from each other. Multiple *Drosophila* species were sampled for DILPs 1-7, which generally clustered into separate “DILP” subcategories, forming monophyletic groups. Only *Drosophila melanogaster* was sampled for DILP8, and this sequence was embedded within DILP2 sequences. DILP groups were interspersed among other ILP samples from other invertebrates, and DILP7 sequences were the most divergent. ILP sequences from invertebrate chordates (lancelets and tunicates) were also dispersed in the tree. Sequences from tunicates (*Ciona*) were most closely related to flatworm sequences. The three lancelet sequences (*Branchiostoma belcheri*) do not form a monophyletic group, with two sequences close to insulin/IGF sequences (see above), and a third sequence (“insulin-like 3”) from this taxon distantly related to the others and embedded in a group of mosquito sequences. All *C. elegans* sequences clustered together with the exception of *C. elegans* peptide 17. The former was most closely related to a mix of sequences from insects, flatworms, and tunicates, while the peptide 17 was most closely related to the DILP4 group. ILP sequences from cone snails (*Conus*) formed a monophyletic group that was divergent from the two other gastropod sequences in the data set (marine sea slug *Aplysia californica* and freshwater snail *Lymnaea stagnalis*).

**Fig 3.**
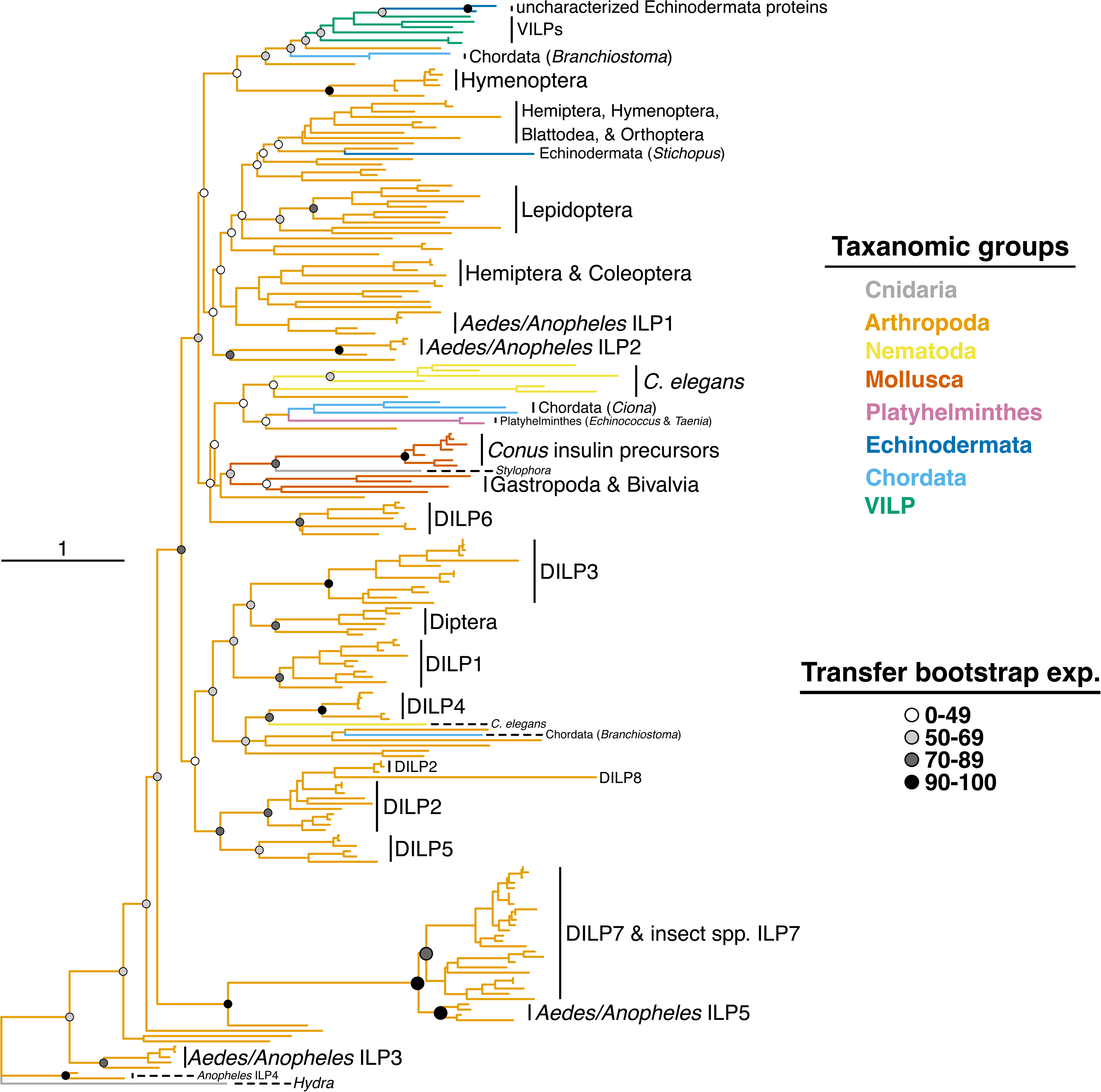
Phylogenetic relationships among invertebrate insulin-like proteins (ILPs) and viral insulin/IGF-1 like peptides (VILPs). The tree was generated using maximum likelihood based on an alignment of 205 sequences. Circles at key nodes illustrate clade support in the form of transfer bootstrap expectation (TBE) values. The removal of vertebrate sequences in this analysis provided a more robust hypothesis of relationships among these sequences and the ability to label more taxa.

The most likely topologies for separate phylogenetic analyses of vertebrate insulin, IGF- 1, and IGF-2 are shown in Fig 4. In each analysis the tree was rooted with the most distantly related sample(s) based on our current knowledge of the vertebrate tree of life. The insulin, IGF- 1, and IGF-2 trees were rooted with hagfish, ray-finned fish, and shark sequences, respectively. Across all three analyses, sequences from the major vertebrate classes typically formed monophyletic groups when multiple species were sampled. For example, in all three analyses the samples of mammals, birds, and ray-finned fishes are most closely related to sequences from their respective taxonomic group. An exception in each analysis is that some reptile sequences are more closely related to sequences from birds than from those of other reptiles, which follows the known finding that Reptilia (as historically defined) is not a monophyletic group [51,52]. The relationships among vertebrate groups generally correspond to expectations based on the current understanding of the tree of life. Notable exceptions include the placement of Chondrichthyes and an Amphibia-Mammalia grouping in the insulin tree, and the placement of three Reptilia sequences as sister to Mammalia in the IGF-1 tree. Each tree in Fig 4 includes a scale bar of 0.3 substitutions/site, and comparing these scales shows that IGF-1 sequences are more conserved among vertebrates compared to insulin and IGF-2 sequences. Also, in each analysis the Actinoptergyii (ray-finned fishes) has the most within-group sequence diversity. For each type of ligand, more detailed figures of relationships among mammal sequences are provided in the supporting information (Fig S1-S3). Generally, sequences from mammalian families [e.g.

**Fig 4.**
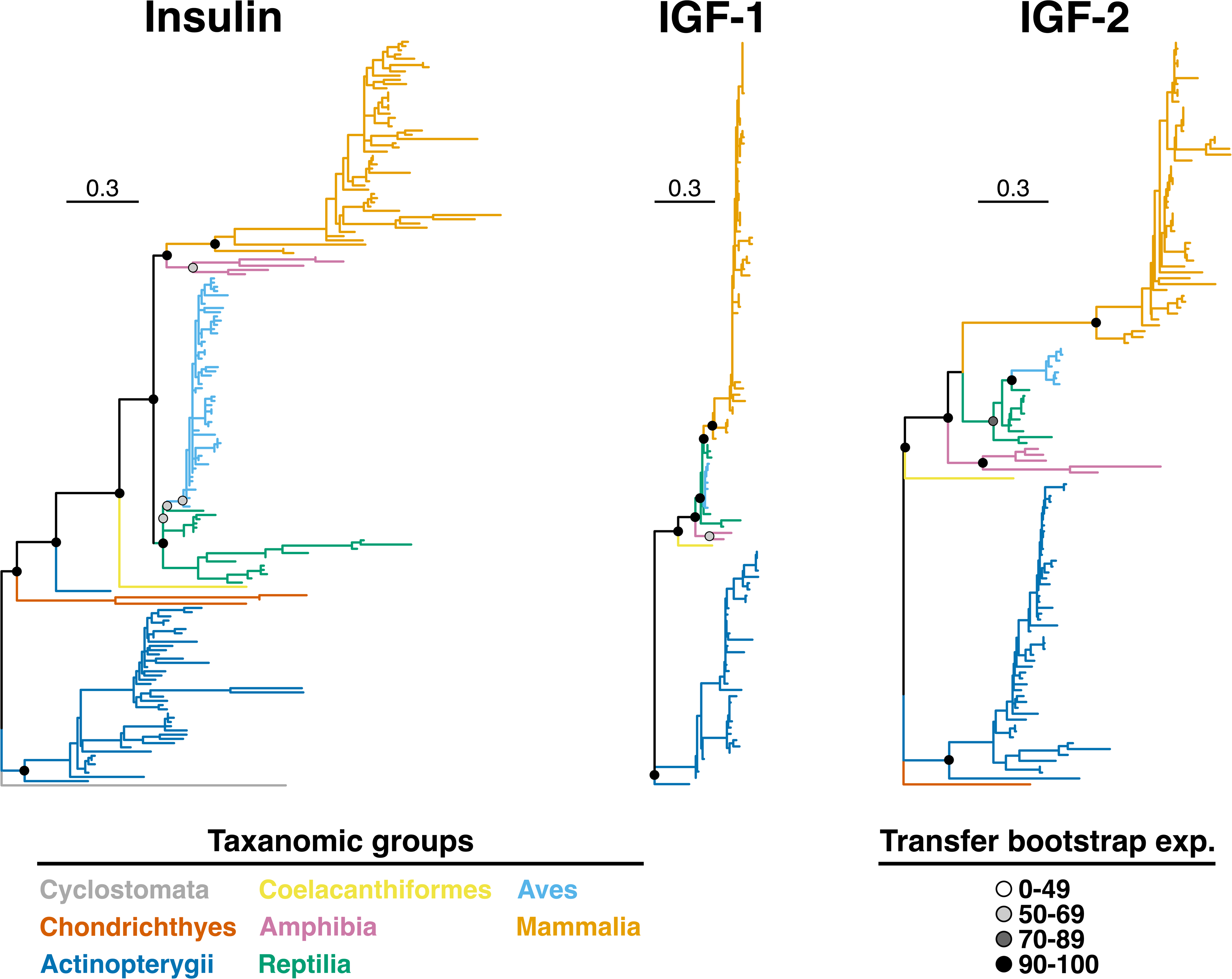
Phylogenetic relationships among vertebrate insulin and insulin growth factor (IGF) ligands. Independent maximum likelihood analyses were conducted for insulin, IGF-1, and IFG- 2 alignments. Circles at key nodes illustrate clade support in the form of transfer bootstrap expectation (TBE) values. The trees are drawn to the same scale, allowing comparison of the diversity and divergence within and among these ligand groups.

Hominidae (great apes), Bovidae (cloven-hoofed ruminants), Felidae (cats)] are more closely related to each other than sequences from other families. As expected from the Mammalia tree of life, sequences from monotremes (duck-billed platypus) and marsupials (e.g., wombat, Tasmanian devil, opossum) are considerably divergent to those from placental mammals.

Moreover, sampled insulin sequences from hystricomorph rodents (guinea pig, degu, mole-rats) were considerably divergent from other rodent sequences.

### Analysis of sequence conservation in vertebrates reveals that IGFs are more conserved than insulin

We created sequence logos for insulin (Fig 5), IGF-1 (Fig 6), and IGF-2 (Fig 7) alignments to analyze sequence conservation within main vertebrate classes. Invertebrate ILPs were not included due to comparatively high sequence diversity. We examined overall conservation of each ligand by determining the percentage of residues fully conserved (i.e., same amino acid in all sequences) across all vertebrates, and those conserved in at least 90% of the sequences within each major vertebrate class (File S2). Throughout this study, sequence positions are assigned based on the analogous human ligand/receptor, with no number assignments for indels introduced into alignments by other species. This analysis revealed that in the insulin B-chain there are eight fully conserved residues (LeuB6, CysB7, GlyB8, LeuB11, ValB12, Cys19, GlyB23, and ProB28), comprising 26% of the human B-chain length. Additionally, there are two (6%) residues (AlaB14 and PheB24) conserved in at least 90% sequences in each class. In the insulin A-chain, only the four diagnostic cysteine residues (19%) are fully conserved (CysA6, CysA7, CysA11, and CysA20), but six other residues (29%) are conserved in at least 90% sequences within each vertebrate class (GlyA1, IleA2, ValA3, LeuA16, TyrA19, and AsnA21) (Fig 5 and File S2). The insulin C-peptide, absent in mature insulin, showed no fully conserved residues and also varied significantly in length among species (Fig S4). However, the pairs of basic residues at the N- (Arg-Arg) and C-terminus (Lys-Arg) required for cleavage of the C-peptide [53] are generally well conserved. The exception is Actinopterygii, where one of the arginines at the N-terminus is missing (Fig S4). Taken together, these results indicate the critical role of the conserved regions in insulin A- and B-chains for its processing, receptor binding, and function.

**Fig 5.**
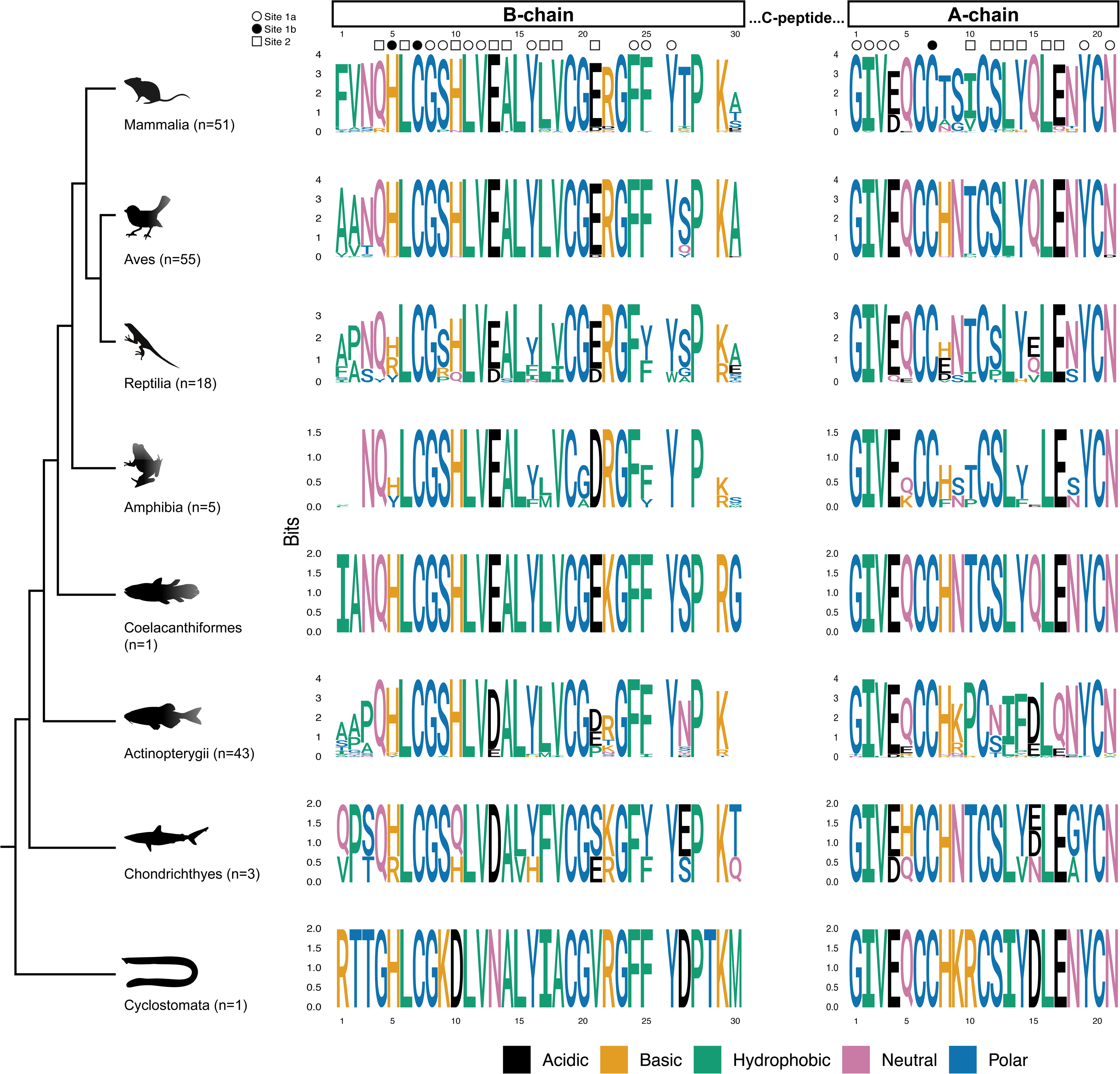
Weblogos of alignments of B- and A-chains of insulin proteins across sampled taxonomic groups of vertebrates. Position numbers are based on the human insulin sequence, and insertions introduced by other taxa are not numbered. Amino acids are color-coded based on their chemistry, and the height of letter corresponds to its amount of information contributed in bits. Proposed binding sites (1a, 1b, and 2; see Introduction) are marked with symbols.

**Fig 6.**
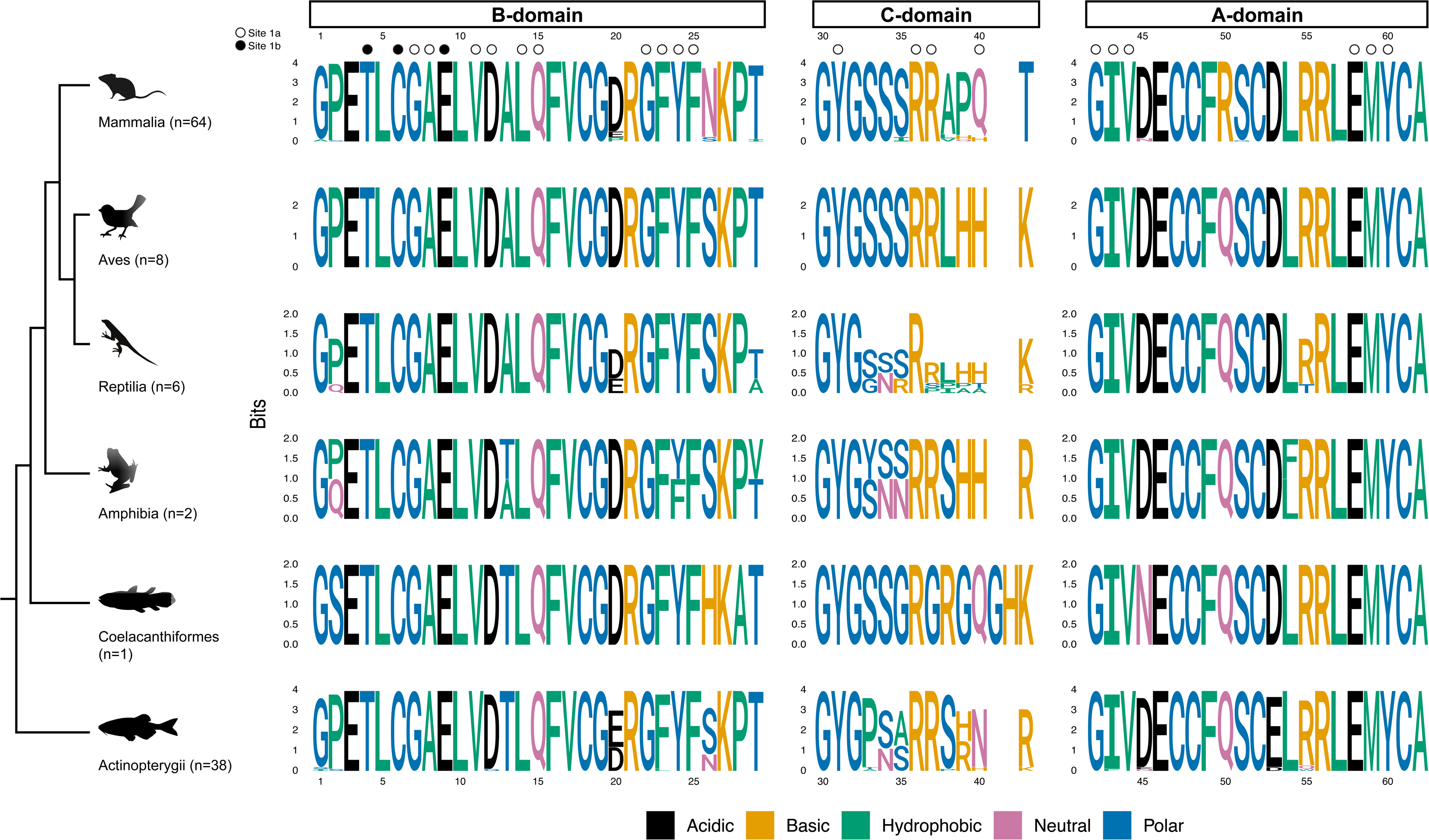
Weblogos of alignments of B-, C-, and A-domains of IGF-1 proteins across sampled taxonomic groups of vertebrates. See Fig 5 caption for more details.

**Fig 7.**
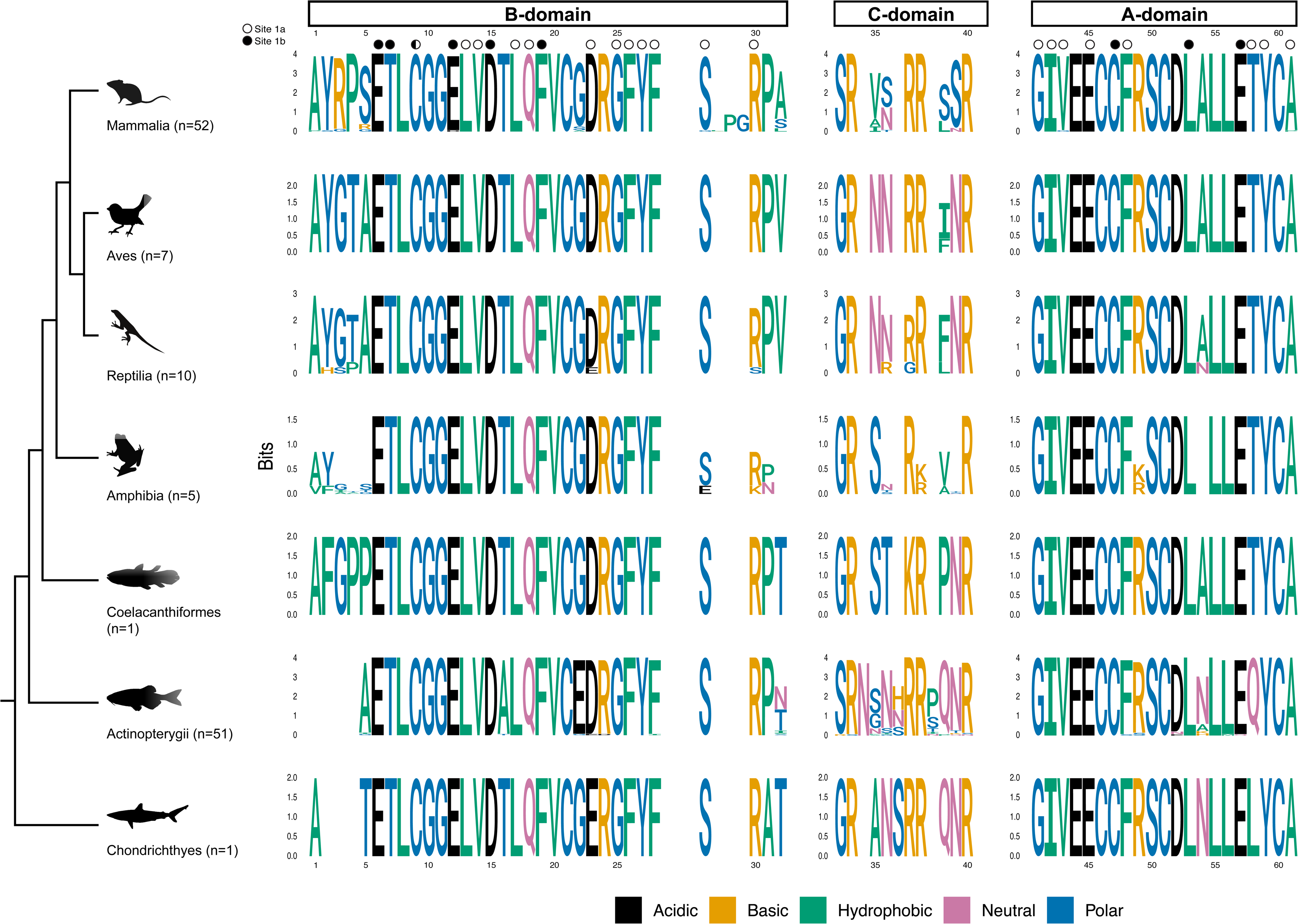
Weblogos of alignments of B-, C-, and A-domains of IGF-2 proteins across sampled taxonomic groups of vertebrates. See Fig 5 caption for more details.

In IGF-1, 20 of 29 amino acids (69%) are fully conserved in the B-domain and 14 of 21 amino acids (67%) are fully conserved in the A-domain (Fig 6 and File S2). Furthermore, 79% and 90% of residues are conserved in at least 90% sequences within each vertebrate class in B- and A-domains, respectively. Positions 2, 13, 20, 24, 26, and 29 in the B-domain and positions 54 and 55 in the A-domain show lower rates of conservation. The C-domain exhibits a comparatively lower conservation rate, with only 33% of residues being fully conserved (Fig 6 and File S2). Analysis of the IGF-2 alignment showed similar patterns of sequence conservation. There was 100% conservation in 17 of 32 amino acids (53%) in the B-domain and 13 of 21 (62%) in the A-domain being fully conserved (Fig 7 and File S2). Further, 72% of the residues in the B-domain and 90% in the A-domain are conserved in at least 90% of sequences within each vertebrate class. The conservation rate in the C-domain of IGF-2 is notably lower, with no residues conserved in all sequences and 3 of 8 (38%) conserved in at least 90% of the sequences within each vertebrate class (Fig 7 and File S2).

Taken together and consistent with the length of the branches in phylogenetic analyses, IGF-1 and IGF-2 are overall more conserved ligands compared to insulin. Moreover, the B- and C-domains of IGF-1 are better conserved compared to these domains in IGF-2. In all three ligands the C-chain/domain has comparatively lower sequence conservation. Consequently, IGF- 1 is the most conserved ligand, followed by IGF-2 and insulin, respectively.

### IGF-1 C-domain shows higher variability in IGF1R binding residues compared to the A- and B-domains

We then examined the conservation of residues crucial for the binding of human insulin (see Materials and Methods for study references) and IGFs to IR and IGF1R, respectively, across different vertebrate classes. Notably, the residues involved in the binding of IGF-1 to IGF1R are almost completely conserved. In IGF-1 B-domain, 7 of 10 (70%) of Site 1a binding residues are fully conserved and another two (90% in total) are conserved in 90% or more sequences in each major vertebrate class. All three Site 1b binding residues are conserved in over 90% sequences IGF-1 B-domain (Fig 6 and File S2).

There are only Site 1a binding residues in IGF-1 A- and C-domains. Five of six residues (83%) are fully conserved in the A-domain, while two of four residues (50%) are conserved in the C-domain, making it the least conserved IGF-1 domain with respect to IGF1R binding residues. The non-conserved binding residue in the A-domain is at the position 43, which is not fully conserved in Actinopterygii. The two non-conserved binding residues in the C-domain are at positions 37 and 40. In position 37, most of the species hold Arg, except for some reptiles (Ser/Pro substitution) and the only Coelacanthiformes sequence in the dataset (Gly substitution). Position 40 shows higher variability with Gln being the predominant residue in mammals (94% sequences) and in the only Coelacanthiformes sequence, while His is the predominant in Amphibia, Reptilia and Aves and Asn. Most Actinopterygii lack this residue, but Asn is predominant in the rest (Fig 6 and File S2).

### Residues important for binding of IGF-2 to IGF1R are well conserved

In IGF-2, we analyzed residues known to be involved in binding to Site1a and Site1b of IGF1R. In the A-domain, 4 of 8 Site 1a residues (Gly41, Ile 42, Glu45 and Y59; 50%) are fully conserved, while three more (Val43, Phe48 and and Ala61; 88% in total) are conserved in at least 90% sequences within each vertebrate class. The remaining Site 1a residue at position 58 is fully conserved within most vertebrate classes, except for Actinopterygii (substituted with Gln) and Chondrychthies (substituted with Leu). Two of three Site 1b residues (Cys47 and Leu53; 67%) are fully conserved in the A-domain (Fig 7 and File S2). In the B-domain, 8 of 12 Site 1a residues (67%) are fully conserved and one more (75% in total) is conserved in at least 90% sequences in each vertebrate class. Moreover, 5 of 6 (83%) Site 1b residues are fully conserved in the B-domain of IGF-2, with the remaining one being conserved in at least 90% sequences in each vertebrate class. No IGF1R binding residues were identified in the C-domain of IGF-2, although Arg38 may potentially form a stabilizing salt bridge with human IGF1R residue Glu305 [54]. Arg38 is fully conserved in all major vertebrate classes, except for Amphibia (substituted with Lys in 60% of sequences).

### Insulin residues interacting with IR binding Site 1 are more conserved than residues interacting with IR binding Site 2

Compared to IGF-1 and IGF-2, we observed more variation in the insulin residues binding to IR. In the insulin B-chain, 3 of 8 (38%) of Site 1a residues and 1 of 2 (50%) of Site 1b residues are fully conserved. In addition to that, one more Site 1a residue (50% in total) is conserved in at least 90% sequences in each class. Interestingly, in the A-chain, no Site 1a residue is fully conserved, but 5 of 6 (83%) of Site 1a residues are conserved in at least 90% sequences in each class. This is due to numerous but low frequency polymorphisms (Fig 5, Fig S4, and File S2).

The only Site 1b residue in insulin A-chain is CysA7, which is fully conserved in all sequences due to its role in disulphide bond formation. Among the Site 1 binding residues that show lower rates of conservation are positions B16, B25, B26 and A4 (Site 1a) and B5 (Site 1b).

Nevertheless, some of the mentioned positions hold substitutions that are conservative, such as Phe/Tyr fluctuation in position B25 or some Asp substitutions of the most prevalent Glu in position A4.

On the other hand, only 1 of 8 (13%) of residues are fully conserved in Site 2 in B-chain, with 25% residues being conserved in at least 90% sequences in each class. No Site 2 residues are fully conserved in the A-chain and only one out of 6 (position A16, 17%) is conserved in 90% or more sequences in each class. Positions B10, B13, B17, B18, B21 and A10, A12, A13, A14 and A17 all show lower conservation rates. Notably, at position A10, in which Thr is the predominant residue in most classes, 98% of Actinopterygii have Pro while mammals have Ile/Val. In position A12, Ser is the most prevalent residue in most classes, but Asn predominates in Actinopterygii. Other Site 2 positions with some notable, although predominantly conservative substitutions include A14 (mostly Tyr in all species but mostly Phe in Actinopterygii) and A17 (mostly Glu in all species but mostly Asp in Actinopterygii). Taken together, these results suggest that insulin binding Site 1 is more conserved during the evolution compared to Site 2.

### Residues important for insulin dimerization and hexamerization are generally well conserved

Although insulin is active in its monomeric form, unlike IGFs, it is initially stored in hexameric structures crucial for its biosynthesis, storage, and release. Human insulin forms dimers at micromolar concentrations that further form hexamers in the presence of zinc or other bivalent ions [55]. The residues responsible for insulin dimerization are primarily located in the C-terminus of the B-chain, overlapping with Site 1a binding residues. Specifically, PheB24 and PheB26 play pivotal roles in the dimer formation [55,56]. PheB24 is conserved across all species in our dataset, except for the mammal common degu (*Octodon degus*) where the corresponding residue is missing. Similarly, while most sequences retain Tyr in position B26, three reptiles (all in genus *Gekko*) exhibit Trp substitution in this position. Additionally, ValB12, GlyB23 and ProB28, which are all involved in human insulin dimerization [56], are fully conserved, whereas TyrB16 shows variability across species. While fully conserved in Mammalia, Aves, Coelacanhiformes (n=1) and Cyclostomata (n=1), TyrB16 exhibits varying conservation levels in Actinopterygii (95%), Amphibia (80%), Chondrichthyes (67%) and Reptilia (61%) (Fig 5 and S4).

Three insulin dimers can further form hexamers [55]. In this process, six GluB13 side chains are brought into close proximity, creating mutual repulsion due to their negative charge. Therefore, zinc coordination with HisB10 is crucial for stabilization of the hexamer structure. The destabilizing GluB13 is known to be conserved in all hexamer forming insulins and it was shown previously that its mutation to charge free Gln leads to stable zinc free insulin [57]. While HisB10 is conserved in most species, exceptions include two mammals (*Octodon degus* and *Cavia porcellus*) and one avian species (*Buceros rhinoceros*), where it is substituted with Asn.

Some reptilian sequences (genus *Gekko*) show a Gln substitution for HisB10. Only one sequence out of three Chondrichthyes in the dataset preserves His in the position B10, while the other two hold Gln in the position. The only Cyclostomata sequence in our dataset carries Asp in the position B10. On the other hand, GluB13 is fully conserved in mammals, birds, amphibians, and Coelacanthiformes (n=1), while some reptiles and fish species show substitutions with Asp, preserving the negative charge. The only Cyclostomata sequence in the dataset carries Asn in B10 position, suggesting the loss of the negative repulsion upon insulin hexamer formation (Fig. 5 and S4).

### Viral insulins contain 13 amino acids that are well conserved across species in insulin and IGFs

VILPs were recently discovered and characterized [35,45–48] and because their natural processing remains unknown, we chemically synthesized VILPs based on their alignment with human insulin and IGFs. These VILPs have been shown to be active [35,45,47,48], and vary in length from 58 to 64 amino acids. 13 of these amino acids are conserved in all six VILPs (Fig S5). These include all the six cysteines that are conserved within the insulin/IGF-like peptide family (Fig 1A). Other fully conserved residues in the B-chain/domain are LeuB11, AlaB14, LeuB15, GlyB23, TyrB26 and ProB28. The only conserved A-chain/domain residue other than the cysteines is LeuA18. There are no conserved amino acids in the C-chain/domain, which generally shows lowest conservation within VILPs, while the B-chain/domain is the most conserved one. All the residues that are conserved in VILPs are generally well conserved across animal species in all insulin, IGF-1, and IGF-2 (Figs 5-7 and File S2). Exceptions are represented by positions B14, B26 and B28. Regarding the position B14, as in VILPs, Ala is the predominant residue in insulins (position B14) and IGF-1s (position 13, only in amphibia there is equal occurrence of Ala and Thr in this position), but not in IGF-2s, which has Thr well conserved among species in the corresponding position 16 (File S2). TyrB26 is well conserved in insulins (position B26), but not in IGF-1s (fully conserved Phe25) or IGF-2s (well conserved Phe28).

Similarly, Pro in VILP position B28 is fully conserved in insulins (position B28), but not in IGF- 1s (fully conserved Lys27) and IGF-2s (well conserved Arg30). These results indicate that the residues that are conserved in all VILPs share higher similarity with insulin than IGFs, however, it has to be noted that in the case of positions B26 and B28, the VILP corresponding amino acids are well conserved in neighboring positions in both IGFs (Fig. 1A, S6, S7).

Six of the 13 residues conserved in all VILPs are implicated in the interaction of insulin to IR (CysB7, LeuB11, AlaB14, TyrB26, CysA7, and LeuA18). Additionally, four residues (Cys6, Leu14, Gly22 and Tyr24) are involved in binding of IGF-1 to IGF1R.

### Phylogenetic relationships among receptor sequences show that vertebrate IRs and IGF1Rs arose early vertebrate evolution

The evolutionary history of IR and IGF1R sequences was estimated using a multiple sequence alignment of 208 vertebrate IR, 160 vertebrate IGF1R, and 39 invertebrate receptor sequences. The maximum likelihood topology of all receptors was rooted with the fresh-water polyp *Hydra vulgaris* (phylum Cnidaria), the most divergent species in our dataset (Fig 8). Early-branching clades in this topology contain sequences from invertebrates. Vertebrate IR and IGF1R sequences formed reciprocally monophyletic groups, indicating that these receptors evolved from a common ancestor during early vertebrate evolution, possibly from a duplication event. In general, the relationships among major vertebrate groups within both IR and IGF1R clades matched the modern understanding of the vertebrate tree of life [58]. Within the IR and IGF1R phylogenetic groups, Chondrichthyes, Actinopterygii, Aves, and Mammalia were all monophyletic. Within both IR and IGF1R, some sequences from reptiles were more closely related to sequences from birds than from those of other reptiles, in line with the finding that reptiles are not a monophyletic group [51,52]. For amphibians, both IR and IGF1R sequences from the African clawed frog (*Xenopus laevis*) were divergent from sequences of other amphibians and sister to all other tetrapod sequences.

**Fig 8.**
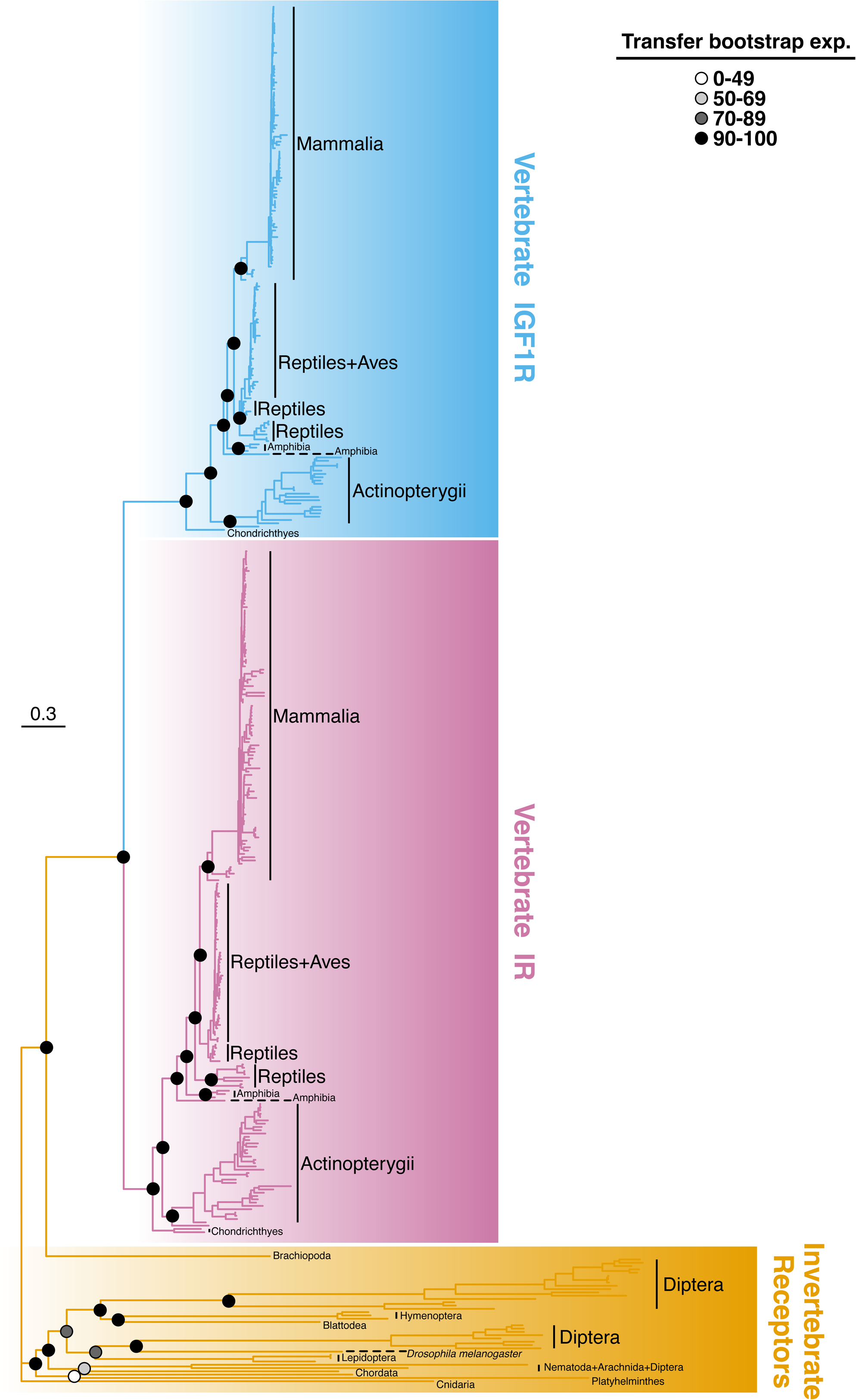
Phylogenetic relationships among vertebrate insulin receptor (IR), vertebrate IGF-1 receptor (IGF1R), and invertebrate receptors. The tree was generated using maximum likelihood based on an alignment of 407 full length sequences. Branches and clades are color-coded for different categories of receptors, and circles at key nodes illustrate clade support in the form of transfer bootstrap expectation (TBE) values.

Previous research on IR and IGF1R evolution focused on full receptor sequences or ectodomains [59,60]. In this study, we analyzed the α- and β-subunits of both vertebrate IR and IGFR independently to identify possible differences in phylogenetic relationships between the ligand binding α-subunits and catalytically active β-subunits. The α-subunits of IR and IGF1R receptors each formed a monophyletic group (Fig S6), and the relationships between vertebrate groups were similar to what was recovered with full length sequences. Similar results were produced using sequences from β-subunits (Fig S7). Taken together, no significant evolutionary differences were observed between the full-length receptors, α-subunits, and β-subunits.

### Analysis of receptor domains involved in ligand binding

To examine the evolution of the receptor domains that are known to be involved in the ligand-receptor interaction of hIR (L1, FnIII-1 and α-CT region) and hIGF1R (L1, FnIII-1, α-CT region and CR), we created sequence logos of vertebrate sequences (Figs 9-11) and calculated conservation scores (File S2) focusing on these domains. In the L1 domains of both IR and IGF1R, most residues are highly conserved within Mammalia and Aves, with greater diversity observed in Reptilia, Amphibia (n=4) and Actinopterygii. A notable exception within mammalian IGF1R is represented by some cetaceans (dolphins and whales), that possess a 15- residue long insertion between residues 1 and 2 (numbered as in human receptor), and the N-terminal regions in mammalian and avian IRs. Amphibian and Actinopterigiian IGF1Rs contain a 2 and 5-7 amino acid long insertion, respectively, between residues 35 and 36, compared to the other classes. Interestingly, in both receptors, some mutations specific for Mammalia were observed. For instance, there is mostly Gly in IR position 10 in mammals, whereas the same position holds mostly Ser in the other vertebrate classes. See IR positions 21-22, 27, 42, 48, 55, 85-86, 104, 116, 147 or 152, and IGF1R positions 6, 13-15, 30, 53, 67, 79-80, and 117 for additional examples (Fig 9 and File S2). Interestingly, a small group of non-placental mammals, including four marsupials (common wombat, Tasmanian devil, koala, and gray short-tailed opossum) and the monotreme platypus, exhibit residues typically found in Aves and Reptilia rather than Mammalia [e.g., IR positions 10 (Ser), 22 (Met), 42 (Lys) or IGF1R positions 53 (Asp), 67 (Ser) and others]. These results suggest that these non-placental mammals have a more ancestral version of IR and IGF1R L1 domains compared to other mammals.

**Fig 9.**
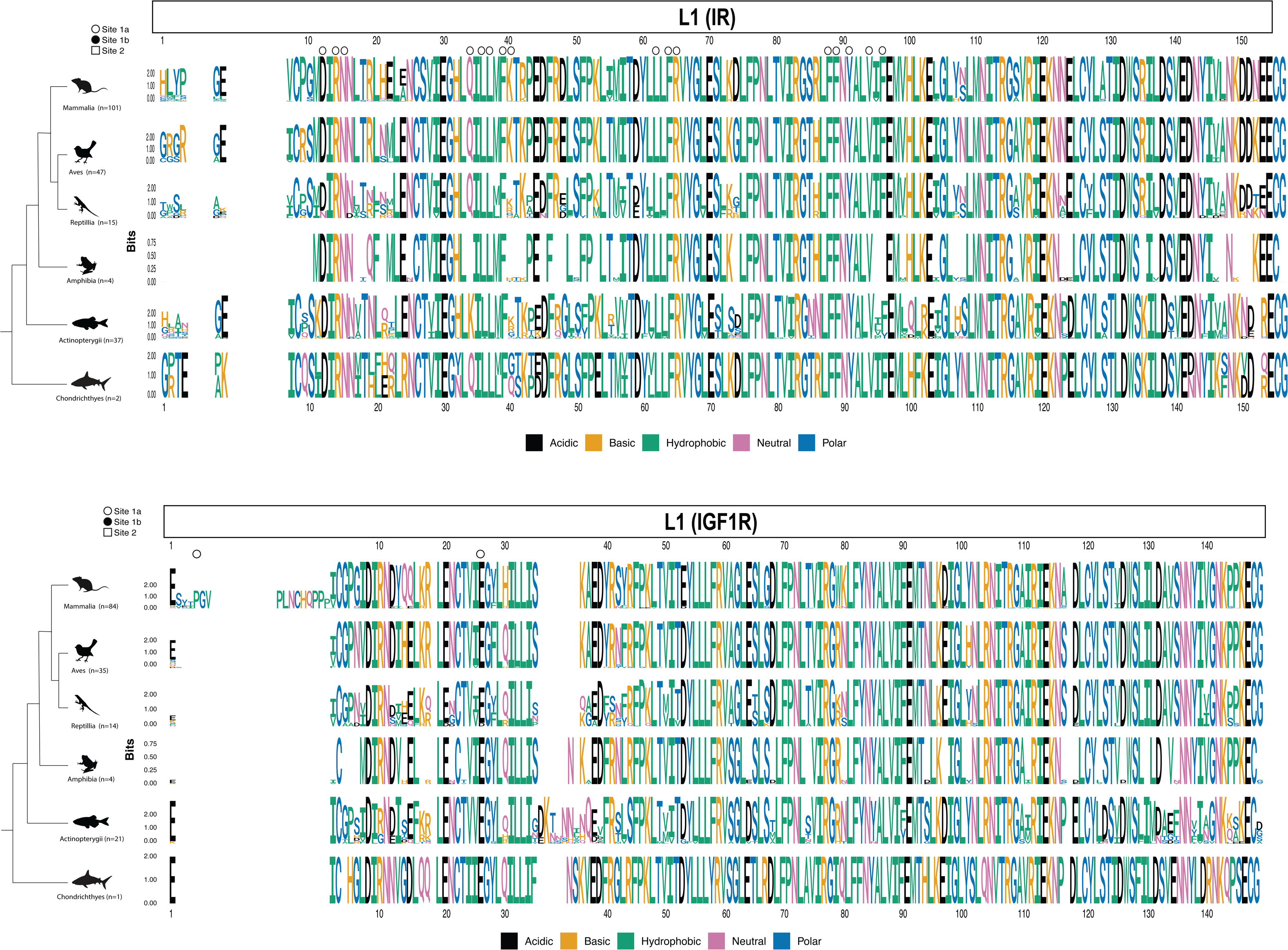
Weblogos of alignments for the L1 domain of sampled insulin receptor (IR) and IGF-1 receptor (IGF1R) sequences. See Fig 5 caption for more details.

The FnIII-1 domain of IGF1R is highly conserved in Mammalia and Aves, with greater diversity observed in Reptilia and particularly in Amphibia and Actinopterygii. The FnIII-1 domain of IR is highly conserved within Aves, but higher diversity can be observed within Mammalia compared to the IGF1R. The highest diversity within Mammalia was observed in the N-terminal region and in the region spanning residues 538-548. Besides high mutation rate, this region is also characterized by sequence length variation among species. Importantly, the diversity of this region is not limited to mammals, but can be noticed in all the vertebrate classes, making it the most diverse region of the IR FnIII-1 domain. Similar to IGF1R, The IR FnIII-1 domains of Reptilia, Amphibia and especially Actinopterygii show higher diversity when compared to Aves and Mammals (Fig 10 and File S2). Generally, this domain seems to be slightly better conserved in the case of IGF1R than IR. An example of the evolutionary trends in IR being influenced by taxonomy is seen in position 538 in mammals. The most prevalent residue, Leu, is found in a variety of orders, including most primates and carnivores, and some rodents and ungulates. Gln predominates in most rodents (i.e., squirrels, mice, rats, hamster), while Met is characteristic for marine mammals (dolphins, whales) and Thr is present in many bovids (e.g., water buffalo, bison, bull, sheep, goat).

**Fig 10.**
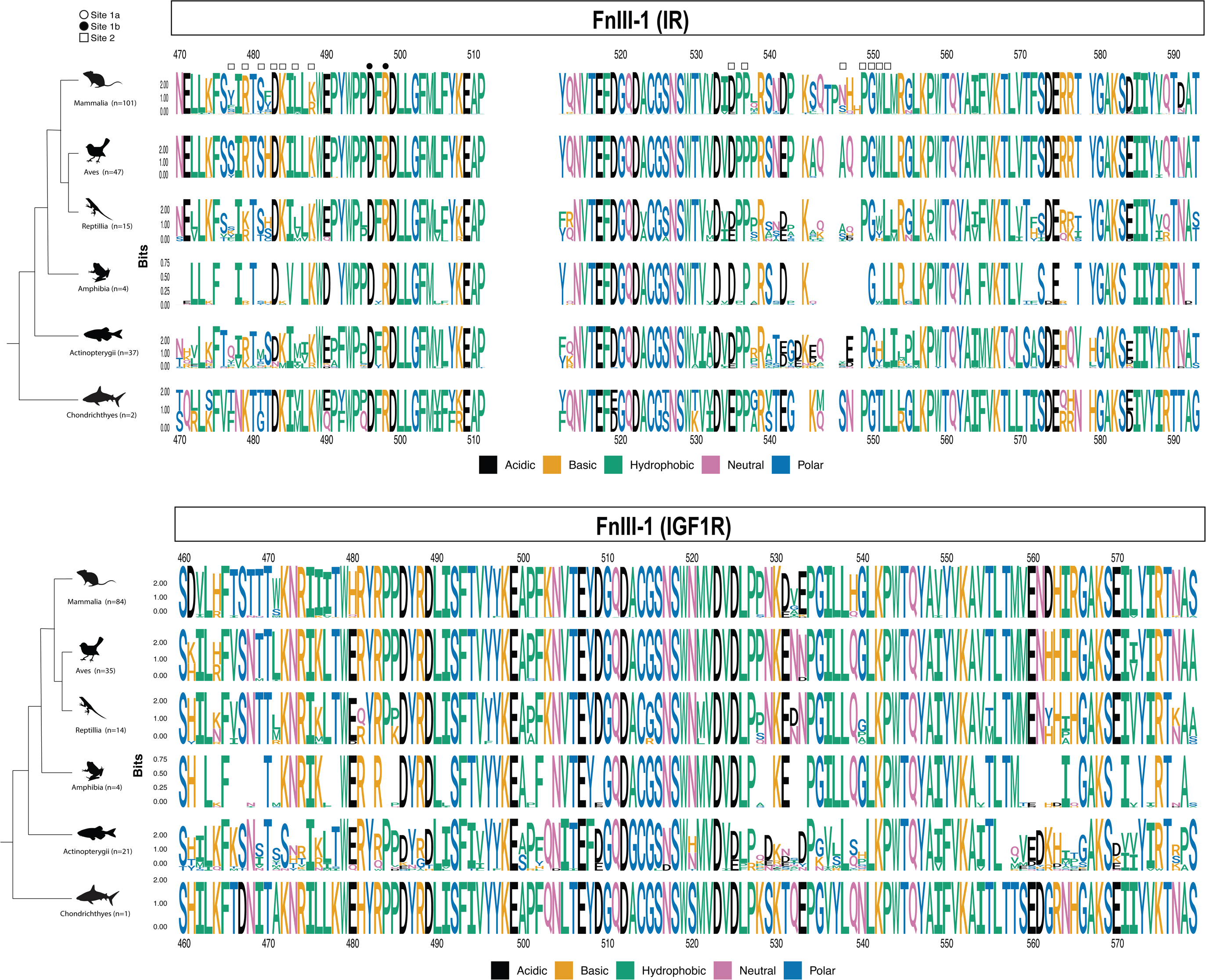
Weblogos of alignments for the FnIII-1 domain of sampled insulin receptor (IR) and IGF-1 receptor (IGF1R) sequences. See Fig 5 caption for more details.

**Fig 11.**
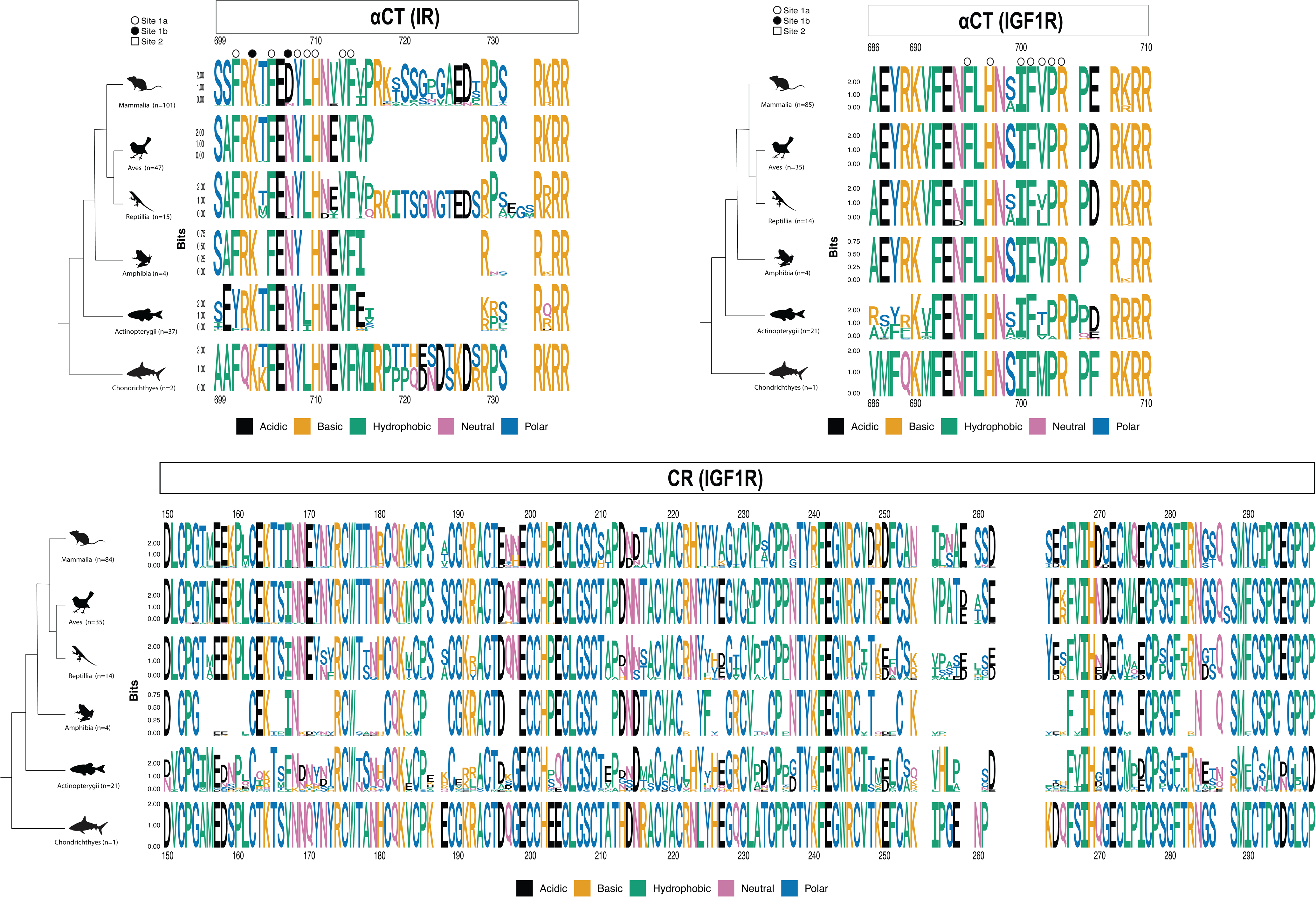
Weblogos of alignments for the α-CT domain of sampled insulin receptor (IR) and IGF-1 receptor (IGF1R) sequences, and the CR domain of IGF1R. See Fig 5 caption for more details.

The IGF1R α-CT region is well conserved across all vertebrate classes. Notable exceptions are represented by the N-terminal region in Actinopterygii, residue 699, that shows Ser/Ala variability in Mammalia, Reptilia, and Actinoprerygii (Fig 11 and File S2). Compared to the IGF1R α-CT region, the IR α-CT shows generally higher variability. Low abundant substitutions are much more frequent than in the case of the IGF1R, and this is in particular true in the case of Mammalia, where only C-terminal residues Arg 732, Arg734 and Arg 735 are conserved in all species in the dataset (Fig S4). A noticeable rate of variability can be observed in the IR-B specific insert (residues 718-729). The existence of alternative splicing in the IR gene, leading to inclusion or exclusion of this insert and therefore generating either IR-B or IR- A, respectively, is known to be specific to mammals. The other vertebrate classes are known to express IR-A-like sequences, but not IR-B [6]. Compared to the rest of amino acids in the mammalian IR α-CT, higher mutation rate was observed in this region. Surprisingly, an analogous insert sequence was identified in IR α-CT positions 717-728 in 40% of the reptilian sequences, specifically turtles and tortoises (File S2). This insert not only shows analogy to some of the most prevalent amino acids of the mammalian IR-B specific insert (Ser721, Gly722, Gly724, Glu726, Asp727), but it is fully conserved within all the reptilian species where it was identified. In addition to this, two Actinopterygii sequences and the two Chondrichthyes sequences in the dataset also contain an insert in the positions 718-729, but there is no significant sequence convergence between these inserts.

Finally, we analyzed sequence conservation in the CR domain of IGF1R (not IR), as this domain is involved in the binding of IGF-1 to the receptor [41,44], but not of insulin to IR [39,40,42,43,61]. The IGF1R CR domain is generally highly conserved within Mammalia and Aves, while Reptilia, Amphibia, and particularly Actinopterygii demonstrate a higher rate of variability. Interestingly, we found several positions with mutations specific to mammals. For example, position 196 predominantly contains Glu in mammals, while Asp is the prevalent residue in all other vertebrate classes. Other examples include positions 166, 180, 197, 210, 227, 240, 248, 253-254, 275, 290, and 292 (Fig 14 and File S2). Even though residues in these positions are generally well conserved within mammals, there are some exceptions. Interestingly and similar to the results of L1 domains, the non-placental mammals (i.e., marsupials and platypus) tend to share the same residue with Aves/Reptilia rather than other mammals. For example, at the aforementioned position 196 the non-placental mammals and Aves/Reptilia share Asp (Glu in other mammals). Other examples include positions 166 (Ser in non-placental mammals and Aves/Reptilia versus Thr in most other mammals), 197 (Gln versus Asn), and 240 (Lys versus Arg). This again shows that non-placental mammals tend to have more ancestral polymorphisms compared to other mammals.

## Discussion

In this study, we examined the phylogenetic relationships among 628 ligand and 407 receptor sequences belonging to the family of insulin-like peptides and their cognate receptors. To the best of our knowledge, this represents the largest study of its kind concerning the phylogenetic analysis of the insulin/IGF system. While potential errors in genome assembly and/or annotation could add some noise to the dataset, the large collection of sequences here provide unprecedented ability to examine the evolution of this ligand-receptor family. Our work also determined the evolutionary position of viral insulins, that were discovered and characterized in the recent years. Additionally, we analyzed the conservation of specific ligand residues to better understand their function and importance across the phylogenetic tree.

The phylogenetic tree of ligands (Fig 2) reveals a clear separation between vertebrate and invertebrate ligands consistent with evolutionary classifications. Insulin, IGF-1, and IGF-2 each form a monophyletic group, with IGF-1 and IGF-2 as sister to each other. These results support an ancient origin in vertebrates, with insulin and the IGFs diverging first, followed by a subsequent divergence of the two IGFs. These observations are consistent with previous research [19,22,29,62]. Earlier research proposed that the IGF precursor genes are homologous to and probably originated from the ancestral insulin gene [62] and that IGFs emerged early in vertebrate evolution [22]. Lancelet (or Amphioxus; *Branchiostoma californiensis*), a primitive cephalochordate and a possible extant relative of the invertebrate progenitor from which the vertebrates emerged, was proposed as a possible transitional form connecting insulin and IGF [22]. Consistent with our results, more recent studies confirm that insulin and IGFs became distinct after vertebrates arose, and that IGF-1 and IGF-2 evolved from a more recent gene duplication during vertebrate evolution, possibly during the transition from agnathans to gnathosomes [24,29]. A 2024 study [19] also confirms the same phylogenetic tree, albeit with a smaller dataset than provided in our study. Similar phylogeny of insulin and IGFs was also proposed based on the locations of the ligands’ genes [63,64]. Therefore, with a more comprehensive dataset, we confirmed the previously proposed representation of the phylogenetic tree of the ligands, and provide the opportunity to explore relationships within groups with more resolution.

Insulin is highly conserved compared to other proteins, with an amino acid substitution rate of about 1×10^-9^ locus/year [65,66]. For example, human and hagfish insulins share 65% amino acid identity and superimposable 3D structures, despite diverging 400 million years ago [24,67]. Insulin B-chain was previously proposed to have higher sequence conservation than the A-chain [19]. In our study, we find nuanced support for this proposal. We observed that the B-chain contains more fully conserved residues, whereas the A-chain has a greater number of residues conserved in over 90% of species in the dataset (File S2). This may reflect different evolutionary pressures and functional constraints between the A-chain and the B-chain. In contrast to functional A- and B-chains, we observed a high sequence variability in the C-peptide (Fig S4). This variability was noted in previous studies [62,66], and it is not surprising since the C-peptide is cleaved out during insulin processing and is therefore not a part of the mature hormone. It was previously reported that besides the six invariant cysteine residues, 10 residues in insulin are fully conserved [68]. These residues include four site 1a interacting residues in the A-chain (GlyA1, IleA2, ValA3, TyrA19), four site 1a interacting residues in the B-chain (GlyB8, LeuB11, ValB12, PheB23), one site 2 interacting residue in the B-chain (LeuB6) and one B-chain residue not directly involved in the receptor binding (GlyB23). Our results generally support the high conservation of these residues. However, we identified a few species in which some of these residues are not fully conserved. Specifically, an Actinopterygii species turbot (*Scophthalmus maximus*) carries mutations in three of these amino acids (GlyA1Thr, ValA3Leu and TyrA19Phe). Another Actinopterygii species, greater amberjack (*Seriola dumeroli*) carries a GlyA1Asn mutation. Additionally, exceptions were found in two mammalian species: an IleA2Val mutation in Nancy Ma’s night monkey (*Aotus nancymaae*) and the absence of PheB24 residue in common degu (*Octodon degus*), as described in the result section. Conversely, we found that ProB28 is also fully conserved in our dataset while it was not being reported as conserved previously. Closer examination revealed that this difference was because the species carrying mutations in ProB28 (amphibian *Siren intermedia,* two Actinopterygii species *Platichthys flesus* and *Opsanus tau*, and lamprey *Geotria australis*) [68] are not present in our dataset due to searching/filtering results. Overall, while we confirm the high conservation of the previously reported insulin residues, we also highlight some exceptions identified in our larger dataset. On the other hand, identifying one residue as fully conserved while mutations in this site were previously identified, underscores potential limitations associated with scraping sequences from large databanks.

Another noteworthy observation is the high conservation of avian insulins, particularly the A-chain (Fig 5 and File S2). Birds maintain high plasma glucose levels compared to other vertebrates and exhibit resistance to insulin-mediated glucose uptake into tissues [69–71].

Interestingly, chicken and turkey insulins are 2-3 times more potent than porcine insulin [66,72]. It has been suggested that birds may have developed insulin resistance as an adaptation to the low-oxygen environment in the Triassic period [70]. Even though we lack a specific explanation for the high conservation of the insulin in birds, it is possible that avian insulins were subjected to strong selection pressures related to changes in glucose metabolism, potentially contributing to the development of high potency insulins.

Our results suggest that insulin site 1 binding residues are better conserved than site 2 residues. Site 2 has been proposed to serve as the initial point of insulin contact with the IR, and this initial binding is understood to induce conformational changes in both the hormone and the receptor, facilitating subsequent binding of insulin to site 1. This dual interaction ultimately activates the receptor and triggers downstream signaling [36,42,73–77]. Interestingly, no equivalent of site 2 has been identified in IGF1R. Instead, IGF1R activation is mediated solely through IGF-1 binding to site 1 [44,73]. The absence of site 2 in IGF1R may suggest that the site 2 function in the initial hormone recruitment may have evolved during vertebrate evolution, after the evolutionary divergence of IR and IGF1R. This might potentially explain the higher conservation of insulin site 1 residues compared to the site 2 residues.

It was previously observed that ancient chordate insulins, such as those from hagfish (*Myxine glutinosa*) and lamprey (*Lampetra fluviatilis*) carry multiple mutations in site 2 binding residues, leading to reduced affinity for the IR. Our analysis, which includes hagfish insulin, supports these findings. Importantly, site 2 residues in insulin overlap with the hexamer forming surface, so mutations at these sites potentially affect both binding and structure. For example, hagfish insulin lacks HisB10, a residue critical for hexamer formation that is known to form dimers but not hexamers [65,66]. In our study, we also observed mutations in HisB10 in two of three Chondrichthyes species in the dataset (*Rhincodon typus* and *Chiloscyllium punctatum*).

HisB10 is conserved in all other species, except for certain reptiles (genus *Gekko*), one avian specie (*Buceros rhinoceros*) and hystrocomorph rodents (*Octodon degus* and *Cavia porcellus*). In hystricomorphs and the reptilian species, we also identified substitutions in residues associated with the dimer forming surface. Taken together, these observations, along with prior studies, suggest that site 2 binding and hexamer formation represent evolutionary innovations in insulins that emerged after its divergence from IGF<u>s</u>. Over time, some species appear to have lost the ability to form insulin dimers and hexamers, potentially conferring specific biological advantages. For instance, hystricomorph insulins are known to lack the ability to dimerize and hexamerize [65,78–80], and they may have lost site 2 entirely [68]. These insulins exhibit higher mitogenic potency compared to other species and an elevated mutation rate in the A- and B- chains [62,66]. Consequently, it has been proposed that hystricomorph insulins may represent another class of “insulin like growth factors” [80]. Similarly, the loss of dimerization and hexamerization, potentially coupled with the acquisition of other functional adaptations, may represent an evolutionary adaptive process optimizing biological functions and physiological regulation in certain reptiles [81].

We observed a higher conservation rate of IGFs compared to insulin. In particular, IGF-1 shows surprisingly high amino acid conservation, followed by IGF-2 (Figs 2 and 5-7, File S2).

This phenomenon had not been noted until recently [19]. According to the authors this is surprising because, in contrast to insulin, IGFs do not oligomerize and should therefore have fewer constraints on residue conservation. However, they postulated that unlike insulin that circulates in a free form in the blood, the plasma concentration of active IGFs is regulated by their binding to six IGF binding proteins (IGFBPs) [82]. There is speculation that the conservation of IGFs is likely due to constraints imposed by their cognate receptors and their interactions with the IGFBPs [19]. While this study proposed a similar conservation rate of IGF- 1 and IGF-2, we found that IGF-1 is slightly better conserved than IGF-2 (Figs 2 and 6-7, and File S2). This may be linked to the more promiscuous nature of IGF-2. In contrast to IGF-1, IGF- 2 binds to both IGF1R and IR-A [34,38], and also binds to another non-tyrosine kinase receptor known as IGF-2 receptor (IGF2R; or also known as cation-independent mannose-6-phosphate receptor) [83,84]. Although describing the function of this receptor is beyond the scope of this study, it is worth noting that IGF-2 binding to IGF2R is only known in mammals [85], and thus this interaction may not be a significant influence of the evolution of the hormone in other taxonomic groups. Our results demonstrate that the C-domain contributes most to the lower conservation of IGF-2 compared to IGF-1 (Figs 6-7, File S2). The C-domain is known to be detrimental for binding to IR, while it is involved in binding of IGF-1, but not IGF-2 to IGF1R [44,54,86,87]. This may explain why the IGF-2 C-domain is more tolerant to substitutions compared to the IGF-1 C-domain.

One of the main aims of this study was to determine the position of the VILPs within the phylogenetic tree of the ligands. Interestingly, the VILPs did not form a monophyletic group (Fig 2). We found that MFRV, GIV, SGIV and LCDV-1 VILPs are more closely related to IGFs than to insulin. This aligns with our previous research which highlighted more pronounced IGF-like rather than insulin-like properties in these VILPs [35,45–48,88]. These VILP sequences were clustered with two Echinodermata sequences from draft genomes of two sea stars. The placement of these Echinodermata sequences is surprising, as it is far from another representative of this phyla and other invertebrate sequences. These Echinodermata sequences are marked as “predicted” in the genome annotation, and further analysis of their validity is needed. On the other hand, LCDV-Sa and LCDV-4 VILPs appeared to be more closely related to insulin than to IGFs. While LCDV-4 remains uncharacterized, LCDV-Sa did not show a strong inclination toward either insulin-like or IGF-like functional properties [46].

To examine the evolutionary relationships between the receptors, we analyzed a dataset comprising 39 invertebrate receptors, 208 vertebrate IR sequences, and 160 vertebrate IGF1R sequences. Unlike two earlier studies that examined 39 [59] and 58 [60] receptor sequences in total, our study employs the largest dataset to date and thus represents a significant contribution to this field. Our phylogenetic analysis of full-length receptors shows that early branches are associated with invertebrate receptors, but that the invertebrate receptors were not monophyletic (Fig 8); a sequence from the phyla Brachiopoda was most closely related to vertebrate sequences. The vertebrate IR and vertebrate IGF1R sequences form sister monophyletic groups (Fig 8), consistent with a duplication of a proto-receptor early in the evolution of vertebrates [59]. As expected, invertebrate receptors exhibit highly divergent and elongated branches, while vertebrate receptors are much more conserved. These results are consistent with the previous studies, which analyzed either full-length receptors [59] or receptor ectodomains [60]. To further investigate the phylogeny of the receptors, we conducted separate phylogenetic analyses of receptor α- (Fig S6) and β-subunits (Fig S7) to assess whether either the function of ligand binding (α-subunit) or the catalytic tyrosine kinase activity (β-subunits) may play specific roles in receptor evolution. To our knowledge, this type of comparison has not been reported previously. Interestingly, we did not observe any significant differences between the phylogenies of the individual subunits and the full-length receptors. Moreover, branch length comparisons indicate that the overall conservation of the α- and β-subunits is comparable. This suggests that both the extracellular ligand binding domain and the intracellular catalytically active domain are subject to comparable selection pressures.

To better understand the conservation of the domains involved in ligand binding in both IR and IGF1R, we conducted a comprehensive analysis. One of the interesting observations was that non-placental mammals (marsupials and platypus) exhibit mutations in the L1 domain of IR, L1 domain of IGF1R, and CR domain of IGF1R that are more typical of Aves and Reptilia than placental mammals. This observation is consistent with a more derived function of these domains in placental mammals.

One of the most interesting findings of this study was the identification of the IR-B specific 12 amino acid insert in the α-CT domain in non-mammals, particularly in some reptiles (turtles and tortoises). This is a significant observation because the presence of this insert (exon 11) in the IR gene has previously been reported to be limited to mammals, where it is alternatively spliced [5,59] to generate more metabolic IR-B and more mitogenic IR-A isoforms. Therefore, the identification of a sequence with homology to the mammalian IR-B specific insert in reptiles is novel and has not been reported before. Whether this sequence also undergoes alternative splicing to produce different receptor isoforms and what evolutionary drivers and functional outcomes are associated with this insert in turtles and tortoises would require further investigation.

The conservation of sites in IR and IGF1R was previously mapped by Rentería et al. [60], which identified a conserved surface in the L1 domain of IR (positions 12, 14-15, 34, 36-37, 39, 59-60, 62, 64-65, 67, 87-91, 94, 96-97, 114, 118, and 120-121) containing residues previously shown to be critical for the ligand-receptor interaction in mutagenesis studies [89–94]. Here, we observe that most of these positions were conserved across most species, but with increased sampling we did identify some low frequency polymorphisms (Fig 9 and File S2). For example, Leu36 (Ile substitution in one Actinopterygii), Tyr60 (Phe substitution in five Actinopterygii) and Lys121 (Arg substitution in one Mammalia and Glu substitution in one Actinopterygii).

Similar to IR, IGF1R has been shown to have conserved regions in the L1 (residues 5, 8, 11, 28, 30, 32-33, 54, 56, 58-59, 79, 81-83, 85, 88, 90, and 112) and CR (residues 240-242 and 252) domains [60], which is supported by previous mutagenesis studies [95,96]. Compared to IR, we observed much higher variation in these residues (Fig 9 and File S2). None of the listed residues were conserved across all species in our data set, with residues 5, 8, 11, 28, 30 and 79 (L1 domain) and 251 (CR domain) showing the highest variability. Lastly, Rentería et al. [60] found positions in the IR and IGF1R α-CT peptide that were fully conserved (positions 702-706, 708- 711, and 713-714). Within mammals, we found that some amino acids spanning this region are missing in two primates (pygmy chimpanzee and silvery gibbon), and that marsupials had a Thr to Met mutation in position 704. Besides mammals, some variability was also observed in position 702 (Actinopterygii), 709 (50% of Amphibia and some Actinopterygii), and 711 (Reptilia). Similarly, Rentería et al. [60] observed almost full conservation of the IGF1R α-CT peptide (residues 686-704). We confirm their findings that residues 692-689, as well as 690, 701 and 704, are fully conserved across species. However, residues 686, 687, 691, 699 and 702 were reported to show some variability and our results support this observation. Unlike in the previous study, we observed that residues 688 and 689 are not fully conserved, with substitutions found in 48% of Actinopterygii.

In conclusion, this study provides a comprehensive analysis of the evolutionary relationships within the insulin/IGF family, leveraging the largest dataset of ligands and receptors to date. Our results confirm previous phylogenetic classifications while offering new insights into the conservation and variability of key residues in both insulin and IGF ligands and their receptors. Notably, we observed exceptions to previously reported conservation patterns, including variations in certain residues across species and new findings regarding the evolutionary divergence of viral insulins and receptor subunits. These findings enhance our understanding of the molecular evolution of the insulin/IGF system and underscore the complexity of evolutionary pressures shaping ligand-receptor interactions across different species.

## Materials and Methods

### Sequence collection and filtering

To collect insulin, IGF, and invertebrate ILP sequences, we screened the protein databases of NCBI and UniProt for entries containing either “insulin” or “IGF*” in the accession name. The resulting sequences were filtered to remove entries that included the words “receptor,” “partial,” “predicted,” “flags,” or “unnamed” in the description. Also, if multiple identical sequences from the same species were recovered then only one record was retained. Additional screening and searching was done to increase taxonomic diversity (e.g., add non-insect invertebrates) and ensure that model species (e.g., mouse, pig, zebrafish, fruit fly, *C. elegans*) were represented. All six known VILP sequences were downloaded and added to the dataset. Since the phylogenetic placement of VILPs was an aim of the study, each VILP sequence was compared to the NCBI protein database using blastp [97] and sequences with > 40% similarity were added to the data set. Separate fasta files were created for vertebrate insulin/IGF sequences, as well all invertebrate ILP sequences. Preliminary alignments were generated using MAFFT v7 [98], and sequences that were aberrant compared to others in their taxonomic group, or atypically long in length, were removed or trimmed, respectively.

After data filtering, we retained 177 insulin, 119 IGF-1, 127 IGF-2, 199 ILP, and 6 VILP sequences (*N* = 628) for downstream analyses (Table 1 and File S1). Sampling of ILPs was dominated by Arthropoda insects (168 sequences), and particularly *Drosophila* (70 sequences). Sampled insulin and IGF sequences represented most major vertebrate groups, with the highest representation in Mammalia (167 sequences) and Actinopterygii (ray-finned fishes; 132 sequences).

**Table 1.**
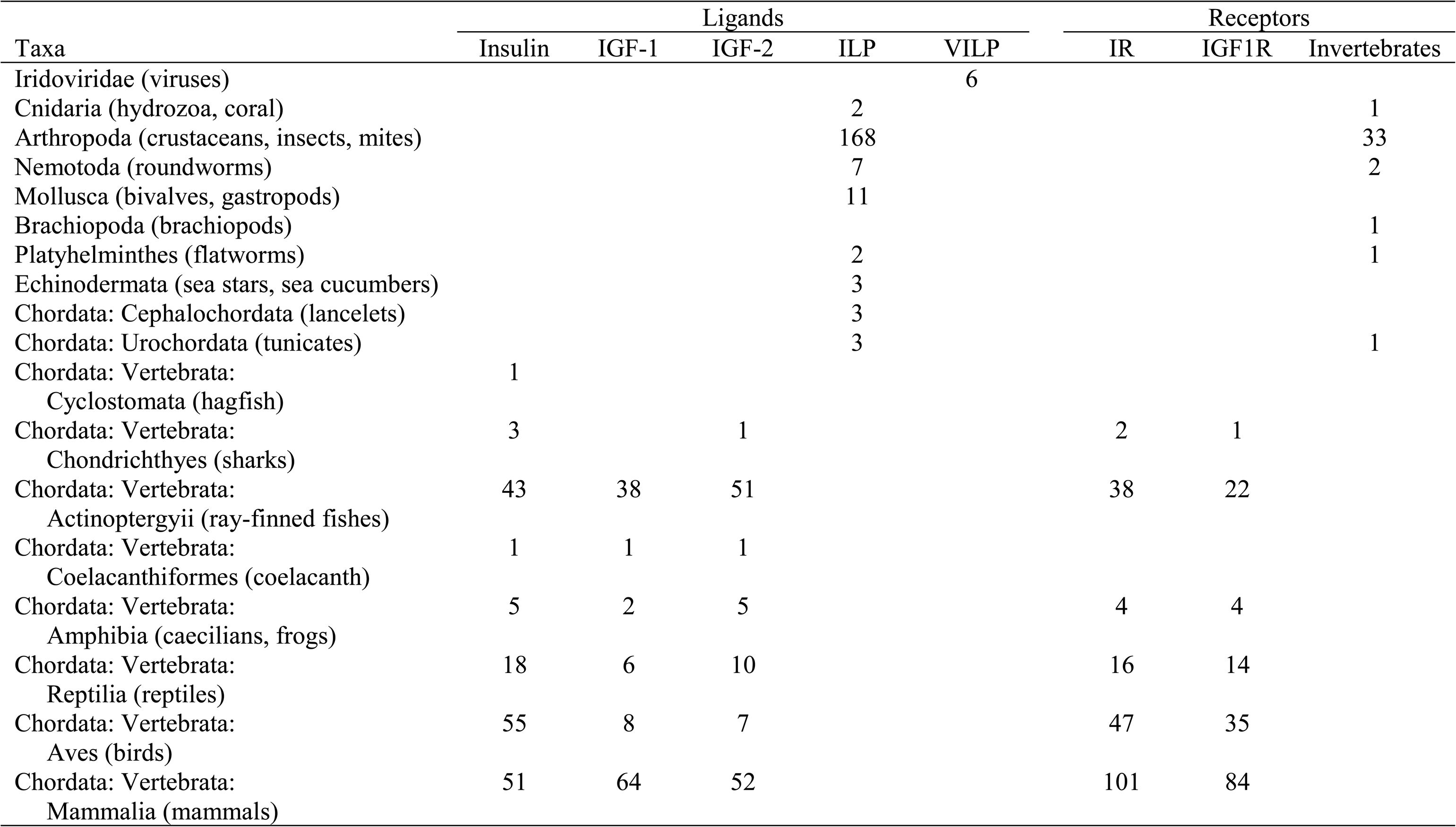
Summary table of ligand and receptor sequences analyzed in this study.

Vertebrate receptor protein sequences were obtained from the NCBI protein database, searching for sequences containing “IGF1R” and “IR” in the accession name. Human IR (isoform IR-B) and IGF1R α- and β-subunits subunit sequences were extracted from Uniprot entries P06213 and P08069, respectively. Receptor subunits (α and β) from other species were then determined by extracting segments that aligned to the human subunits. Subsequent phylogenetic analyses for each data matrix was performed using the methodology described above for ligands. Only Refseq entries were selected, and sequences containing words “partial” or “predicted” were filtered. Initial alignments were performed using MAFFT v7 and aberrant sequences were filtered, yielding 208 IR, 160 IGF1R, and 39 invertebrate receptor sequences **(Table 1)**.

### Alignment and phylogenetic analyses

Estimates of phylogenetic gene trees were generated using an alignment of all ligands, and for individual groups (i.e., insulin, IGF-1, IGF-2, ILPs+VILPs). Multiple sequence alignments were generated using MAFFT to infer sequence homology, and maximum likelihood phylogenies for each data matrix were inferred using RAxML-NG v0.9 [99]. The appropriate model of protein evolution for each analysis was determined with ModelTest-NG v0.1 [100] using the AICc option. Each analysis comprised 20 searches with different starting trees, and retained the tree with the highest likelihood score. Support for relationships in the best tree from each analysis was evaluated with the TBE metric, which has been shown to accurately measure node robustness and works particularly well for deep nodes [101]. The number of TBE replicates was determined with the autoMRE algorithm (default convergence cutoff 0.3), with a maximum of 1000 replicates.

Phylogenetic trees were similarly estimated for alignments with all receptor sequences and each separate group (i.e., IGF1R, IR, IGFR α-subunit, IGFR β-subunit, IR α-subunit and IR β-subunit). Full length receptor alignments were parsed into separate α- and β-subunit matrices using knowledge of human receptors as a reference.

All alignment and tree files produced by this study are available at https://github.com/jeffdacosta/evo-insulin-IGF.

### Motif characterization and conservation

To visualize the conservation of sequences across different taxonomic groups, we generated sequence logos for each alignment using the R package ggseqlogo [102]. Separate sequence logos were generated for biologically relevant subsections (e.g., insulin A- and B-chains) and/or taxonomic groups (e.g., Mammalia, Aves). The boundaries of the chains/peptide/domains were set using the corresponding human sequence.

Sequence conservation among ligands and receptors was compared across vertebrate groups, with particular scrutiny of ligand residues that have been identified as important for IR and IGF1R binding in previous studies [36,39–44,55–57,61,86,103]. Separate conservation calculations were done for each taxonomic group and protein subsection. In each case, we identified the most common amino acid for each position in the alignment, and the percentage of sequences with this most common amino acid. Gaps were treated as a separate character state.

For example, if an alignment had a position with 6 K, 2 R, and 2 gaps then K was designated at the most common amino acid with a conservation percentage of 60.

## Supporting Information

**File S1.** Excel spreadsheet with details on the sampling of insulin/IGF ligand and receptor sequences.

**File S2.** Excel spreadsheet with results of analyses of conservation in insulin/IGF ligand and receptor sequences.

**Fig S1.** Phylogenetic relationships among sampled mammalian insulin sequences.

**Fig S2.** Phylogenetic relationships among sampled mammalian IGF-1 sequences.

**Fig S3.** Phylogenetic relationships among sampled mammalian IGF-2 sequences.

**Fig S4.** Weblogos of alignments of C-chains of insulin proteins across sampled taxonomic groups of vertebrates.

**Fig S5.** Sequence alignment and weblogos of the six viral insulin/IGF-1 like peptides (VILPs) analyzed in this study.

**Fig S6.** Phylogenetic relationships among the α-subunit region of vertebrate insulin receptor (IR) and vertebrate IGF-1 receptor (IGF1R) sequences.

**##Fig S7.** Phylogenetic relationships among the β-subunit region of vertebrate insulin receptor (IR) and vertebrate IGF-1 receptor (IGF1R) sequences.

