## Supporting Figures for "Evolution of Insulin, Insulin-like Growth Factor, and Their Cognate Receptors in Vertebrates, Invertebrates, and Viruses"

**Supporting Information Figures**

**Fig S1.** Phylogenetic relationships among sampled mammalian insulin sequences. This figure zooms in on the Mammalia clade in Fig 4 and labels all tips. The tree was estimated using maximum likelihood on an alignment of 177 vertebrate insulin sequences. Circles at key nodes illustrate clade support in the form of transfer bootstrap expectation (TBE) values.

**Fig S2.** Phylogenetic relationships among sampled mammalian IGF-1 sequences. This figure zooms in on the Mammalia clade in Fig 4 and labels all tips. The tree was estimated using maximum likelihood on an alignment of 119 vertebrate IGF-1 sequences. Circles at key nodes illustrate clade support in the form of transfer bootstrap expectation (TBE) values.

**Fig S3.** Phylogenetic relationships among sampled mammalian IGF-2 sequences. This figure zooms in on the Mammalia clade in Fig 4 and labels all tips. The tree was estimated using maximum likelihood on an alignment of 127 vertebrate IGF-2 sequences. Circles at key nodes illustrate clade support in the form of transfer bootstrap expectation (TBE) values.

**Fig S4.** Weblogos of alignments of C-chains of insulin proteins across sampled taxonomic groups of vertebrates. Position numbers are based on the human insulin sequence, and insertions introduced by other taxa are not numbered. Amino acids are color-coded based on their chemistry, and the height of letter corresponds to its amount of information contributed in bits.

**Fig S5.** Characterization of viral insulin/IGF-1 like peptides (VILPs). (A) Sequence alignment and (B) weblogo plot of six VILP sequences analyzed in this study. In the weblogo, amino acids are color-coded based on their chemistry, and the height of letter corresponds to its amount of information contributed in bits.

**Fig S6.** Phylogenetic relationships among the α-subunit region of vertebrate insulin receptor (IR) and vertebrate IGF-1 receptor (IGF1R) sequences. The tree was generated using maximum likelihood based on an alignment of 366 sequences. Branches and clades are color-coded for different categories of receptors, and circles at key nodes illustrate clade support in the form of transfer bootstrap expectation (TBE) values.

**Fig S7.** Phylogenetic relationships among the β-subunit region of vertebrate insulin receptor (IR) and vertebrate IGF-1 receptor (IGF1R) sequences. The tree was generated using maximum likelihood based on an alignment of 366 sequences. Branches and clades are color-coded for different categories of receptors, and circles at key nodes illustrate clade support in the form of transfer bootstrap expectation (TBE) values.
